# Exploring High-Dimensional Biological Data with Sparse Contrastive Principal Component Analysis

**DOI:** 10.1101/836650

**Authors:** Philippe Boileau, Nima S. Hejazi, Sandrine Dudoit

## Abstract

**Motivation:** Statistical analyses of high-throughput sequencing data have re-shaped the biological sciences. In spite of myriad advances, recovering interpretable biological signal from data corrupted by technical noise remains a prevalent open problem. Several classes of procedures, among them classical dimensionality reduction techniques and others incorporating subject-matter knowledge, have provided effective advances; however, no procedure currently satisfies the dual objectives of recovering stable and relevant features simultaneously.

**Results:** Inspired by recent proposals for making use of control data in the removal of unwanted variation, we propose a variant of principal component analysis, sparse contrastive principal component analysis, that extracts sparse, stable, interpretable, and relevant biological signal. The new methodology is compared to competing dimensionality reduction approaches through a simulation study as well as via analyses of several publicly available protein expression, microarray gene expression, and single-cell transcriptome sequencing datasets.

**Availability:** A free and open-source software implementation of the methodology, the `scPCA R` package, is made available via the Bioconductor Project. Code for all analyses presented in the paper is also available via GitHub.

## 1 Introduction

Principal component analysis (PCA) is a well-known dimensionality reduction technique, widely used for data pre-processing and exploratory data analysis (EDA). Although popular for the interpretability of its results and ease of implementation, PCA’s ability to extract signal from high-dimensional data is demonstrably unstable [31, 15], in that its recovered results can vary widely with perturbations of the data [35]. What is more, PCA is often unable to reduce the dimensionality of the data in a contextually meaningful manner [27, 2]. Consequently, variants of PCA have been developed in attempts to remedy these severe issues, including, among many others, sparse PCA (SPCA) [41], which increases the interpretability and stability of the principal components in high dimensions by sparsifying the loadings, and contrastive PCA (cPCA) [2], which captures relevant information in the data by eliminating technical effects through comparison to a so-called background dataset. While SPCA and cPCA have both individually proven useful in resolving distinct shortcomings of PCA, neither is capable of simultaneously tackling the issues of stability, interpretability, and relevance. We propose a combination of these techniques, *sparse constrastive PCA* (scPCA), which draws on cPCA to remove unwanted technical effects and on SPCA for sparsification of the loadings, thereby extracting stable, interpretable, and relevant uncontaminated signal from high-dimensional biological data.

### 1.1 Motivation

A longstanding problem in genomics and related disciplines centers on teasing out important biological signal from technical noise, i.e., removing *unwanted* variation corresponding to experimental artifacts (e.g., batch effects). A common preliminary approach for accomplishing such a task involves the application of classical PCA to capture and deflate technical noise, followed by traditional statistical inference techniques (e.g., clustering cells, testing for differences in mean gene expression levels between populations of cells) [25]. Such an approach operates under the assumption that meaningful biological signal is not present in the leading principal components (PC), and that the removal of the variance contained therein allows recovery of the signal previously masked by technical noise. Should these assumptions prove unmet, relevant biological signal may be unintentionally discarded, or worse, technical noise may be significantly amplified.

Several more sophisticated approaches have been proposed, including the use of control genes [9, 28] and control samples [10] whose behavior is known *a priori*. Unfortunately, access to such controls may be severely limited in many settings (e.g., as with prohibitively expensive assays). Alternative approaches, for use in settings where control genes or control samples are unavailable, such as surrogate variable analysis [19], reconstruct sources of unwanted variation that may subsequently be controlled for via covariate adjustment in a typical regression modeling framework. In the context of single-cell transcriptome sequencing (scRNA-seq) data, a class of data that has garnered much interest due to the granularity of biological information it encodes, related approaches have been combined as part of the ZINB-WaVE methodology [29], which relies on factor analysis for a zero-inflated negative binomial (ZINB) generalized linear model (GLM) to remove unwanted variation.

Although such approaches have proven useful, model-based techniques rely on assumptions about the data-generating process to target biological signal, warranting that caution be exercised in their use. Additionally, owing to the diversity of experimental settings in high-dimensional biology, such techniques are often targeted to specific experimental paradigms (e.g., bulk RNA-seq but *not* single-cell RNA-seq). Violations of the assumptions embedded in these techniques may often be difficult – impossible, even – to diagnose, leading to a lack of agreement in findings between different model-based approaches when applied to the same datasets. Accordingly, Zhang *et al*. [37] have shown the lack of consensus among model-based differential expression techniques on RNA-seq datasets, demonstrating that their use gives rise to subjective analyses. By contrast, we propose a wholly data-driven approach to removing unwanted variation, harnessing the information contained in control samples, pre-treatment groups, or other signal-free observations, all while enhancing the interpretability and stability of findings by inducing sparsity.

The remainder of the present manuscript is organized as follows. In Section 1.2, contrastive PCA, sparse PCA, and other popular dimensionality reduction techniques are briefly surveyed. Next, in Section 2, scPCA is formally defined and its desirable properties are detailed. A simulation study and several analyses of publicly available microarray gene expression and single-cell transcriptome sequencing data are presented in Section 3 and the analysis of a protein expression dataset is detailed in Section S7, providing a rich comparison of the proposed methodology to other popular techniques currently relied upon for the exploration of high-dimensional biological data. Finally, we conclude by reviewing the effectiveness of scPCA based on the results of these studies and discussing paths for further investigation.

### 1.2 Background

#### 1.2.1 Contrastive PCA

The development of contrastive PCA was motivated by the need to detect and visualize variation in the data deemed most relevant for the scientific question of interest. Given a target dataset believed to contain biological signal of interest and a similar background dataset believed to comprise only noise (i.e., unwanted variation), the cPCA algorithm returns a subspace of the target data that contains (a portion of) the variation absent from the background data [2]. Specifically, cPCA aims to identify the pertinent variation in the target data by *contrasting* the covariance matrix of its features with that of the background data. For example, consider a scRNA-seq dataset whose samples are contaminated by a batch effect. Given a collection of control samples subjected to the same batch effect, cPCA may be used to remove this unwanted technical noise (see Section 3.1).

Algorithmically, cPCA is very similar to PCA. Consider a column-centered target dataset **X**_*n*×*p*_ and a column-centered background dataset **Y**_*m×p*_, where *n* and *m* denote, respectively, the number of target and background observations (e.g., cells) and *p* denotes the number of features (e.g., genes). Denote their empirical covariance matrices by **C_X_** and **C_Y_** and let 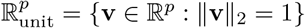 be the set of unit vectors of length *p* (i.e., *p*-dimensional vectors with unit norm). The variances along direction 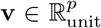 in the target and background datasets are represented by *f*_**X**_(**v**) = **v**^⊤^ **C**_**X**_**v** and *f*_**Y**_(**v**) = **v**^⊤^**C**_**Y**_**v**, respectively. The most contrastive direction 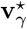 for some fixed contrastive parameter 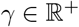 is found by solving

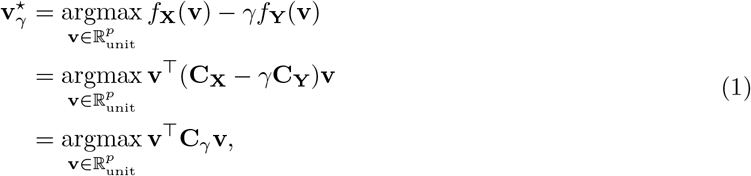

where **C**_*γ*_ = **C_X_** − *γ***C_Y_** is the contrastive covariance matrix [2]. cPCA can therefore be performed by computing the eigenvalue decomposition of **C**_*γ*_. The eigenvectors of **C**_*γ*_ are then used to map the target data to the *contrastive* principal components (cPC).

The contrastive parameter *γ* quantifies the trade-off between each feature’s variances in the target and background datasets. When *γ* = 0, cPCA reduces to PCA – hence, only the target variance *f***_x_**(**v**) is maximized. As *γ* increases, directions that reduce the background variance become more optimal and the cPCs are driven towards the null space of the background covariance matrix. In particular, as *γ* → ∞, the variance in the background data dominates the variance in the target data such that only directions spanned by the background dataset are captured. This is akin to projecting the target data on the null space of the background data and then performing PCA on the projected data [2]. The effect of the contrastive parameter is illustrated in Figure S1 for simulated data similar to those used by Abid *et al*. [2].

Although cPCA offers a novel approach for the removal of unwanted variation, it possesses some drawbacks. In particular, no rigorous framework exists for selecting the contrastive parameter *γ* in order to achieve the optimal amount of contrast between the target and background data. Indeed, Abid *et al*. [2]’s semi-automated approach to selecting an appropriate *γ* relies on visual inspection. Alternatively, recent work by Fujiwara *et al*. [8] introduces a heuristic algorithm based on a one-dimensional rotation of the target data. Additionally, as with PCA, loading vectors may be highly variable and difficult to interpret in high dimensions since they represent linear combinations of all features in the dataset. Relatedly, cPCs are not certifiably free of unwanted technical and biological effects, potentially obscuring relevant biological signal. Unwanted variation can only be removed from the target data if it is also captured by the background data. This issue is only exacerbated as the dimension of the subspace orthogonal to the background data increases, jeopardizing the stability of the cPCs and enfeebling conclusions drawn from them. Work by Severson *et al*. [30] provides a probabilistic framework for linear and non-linear contrastive models that attempts to remedy these shortcomings, though it is accompanied by the typical computational and epistemological issues associated with Bayesian methods.

#### 1.2.2 Sparse PCA

In addition to being difficult to interpret, the PCs generated by applying PCA to high-dimensional data are generally unstable, that is, they are subject to major changes under minor perturbations of the data; we refer to Johnstone and Paul [16] for a recent review. Luckily, an abundance of techniques for sparsifying PCA loadings have been developed to mitigate these issues; we direct the interested reader to Zou and Xue [40] for a recent review. Here, we consider the SPCA technique developed by Zou *et al*. [41]. In contrast to standard PCA, SPCA generates interpretable and stable loadings in high dimensions, with most entries of the loading matrix being zero.

SPCA was born from the geometric interpretation of PCA, which reframes PCA as a regression problem. Given a matrix **V**_*p*×*k*_ whose columns form an orthonormal basis, the objective is to find the projection 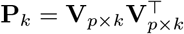 producing the best linear manifold approximation of the data **X**_*n×p*_. This is accomplished by minimizing the mean squared error:

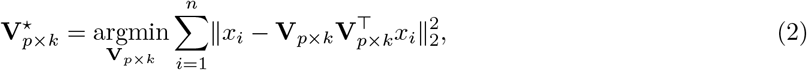

where *x_i_* is the *i*^th^ row of **X** and 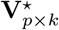 is exactly the loading matrix of the first *k* PCs [40]. A sparse loading matrix can be obtained by imposing an elastic net constraint on a modification of this objective function.

Zou *et al*. [41] show that optimizing the following criterion provides loadings of the first *k* sparse PCs of **X**:

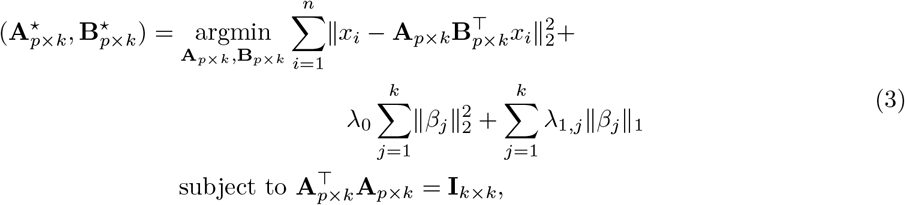

where *β_j_* is the *j*^th^ column of **B**_*p*×*k*_ and where λ_0_ and λ_1,*j*_ are, respectively, the *ℓ*_2_ and *ℓ*_1_ penalty parameters for the non-normalized *j*^th^ loading *β_j_*; as in the original SPCA manuscript the *ℓ*_1_ penalty is allowed to be loading-specific [41]. Then, the sparse loadings are the normalized versions of 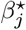, i.e., the vectors 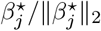.

Zou *et al*. [41] also show that the full dataset need not be used to optimize the criterion; indeed, only the Gram matrix 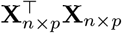 is required. This follows from the first term on the right-hand side of Equation (3), which can be rewritten as 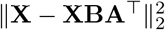. Letting **A**_⊥_ be any orthonormal *p* × (*p* – *k*) matrix such that the horizontal concatenation [**A**; **A**_⊥_]_*p*×*p*_ is orthonormal, we have, from Zou *et al*. [41]’s derivation, that 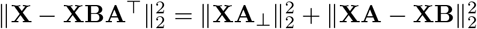. Hence, obtaining the optimal *β_j_* for a fixed **A** is equivalent to minimizing

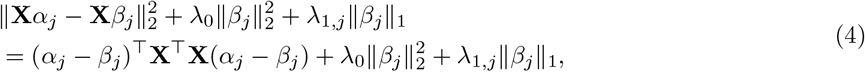

where *α_j_* is the *j*^th^ column of **A**.

Although SPCA provides a transparent and efficient method for the sparsification of PCA’s loading matrices and hence the generation of stable principal components in high dimensions, its development stopped short of providing means by which to identify the most relevant directions of variation in the data, presenting an obstacle to its efficacious use in biological data exploration and analysis. This motivates the development of exploratory methods that build upon the strengths of both SPCA and cPCA.

#### 1.2.3 Other Competing Methods

Other general methods frequently employed to reduce the dimensionality of high-dimensional biological data include t-distributed stochastic neighbor embedding (t-SNE) [32] and uniform manifold approximation and projection (UMAP) [23] (e.g., [3, 4]). Unlike PCA, SPCA, and cPCA, both are nonlinear dimensionality reduction techniques, that is, they do not enforce linear relationships between features. Such a relaxation permits the capturing of local nonlinear structures in the data that would otherwise go unnoticed, though neither approach guarantees that their low-dimensional embeddings reflect the global structure of the data. Becht *et al*. [4] demonstrated the computational efficiency exhibited by these techniques in their application to large datasets, while Amir *et al*. [3] and Becht *et al*. [4] illustrated the stability of their findings, further increasing their popularity as methods of choice for EDA in computational biology. Yet, the flexibility and speed of t-SNE and UMAP come at a cost: (1) these techniques are not endowed with the ease of interpretability of factor analysis methods, (2) the results are sensitive to hyperparameter values (e.g., perplexity for t-SNE and number of nearest neighbors for UMAP) and initializations, (3) identifying reasonable hyperparameters and initializations can prove difficult [17], and (4) the results are inherently stochastic. In lacking an interpretable link between the data’s features and low-dimensional representation, their use as hypothesis-generating tools is restricted. Furthermore, like PCA, neither t-SNE nor UMAP have the ability to explicitly remove unwanted technical effects.

Though the dimensionality reduction methods discussed thus far can be applied to a variety of high-dimensional biological data, still many others have been developed expressly for use with specific high-throughput assay biotechnologies. One such method, ZINB-WaVE, relies on a zero-inflated negative binomial model to better account for the count nature, zero inflation, and over-dispersion of scRNA-seq data, and has been shown to outperform less tailored techniques such as t-SNE [29]. Unlike more general factor analysis methods (e.g., PCA), ZINB-WaVE takes advantage of the rich annotation metadata that are often available with scRNA-seq datasets to remove sources of unwanted variation, while preserving global biological signal [29]. Analogous to PCA, the latent factors produced by ZINB-WaVE are not sparse. Technical and biological noise may remain after taking into account known and unknown sources of unwanted variation [29], potentially blurring any meaningful interpretation of latent factors. Other methods for reducing the dimensionality of scRNA-seq data, such as scVI [21] and SIMLR [33] attempt to identify non-linear latent variables using, respectively, a variational auto-encoder framework and a similarity-learning framework relying on multi-kernel learning. Further discussion of such techniques lies outside the scope of the present work on account of the dissimilarity to methods inspired by factor analysis, like SPCA, cPCA, and ZINB-WaVE, which are our focus.

## 2 Methodology

### 2.1 Sparse Contrastive PCA

The sparse contrastive PCA (scPCA) procedure applies SPCA with minimal modifications to a pair of target and background datasets’ contrastive covariance matrix **C**_*γ*_. The numerical solution to the SPCA criterion of Equation (3) is obtained by the following alternating algorithm until convergence of the sparse loadings [41], where **A**_*p*×*k*_ is initialized as the matrix of loadings corresponding to the *k* leading principal components of **C**_*γ*_:

**For fixed A:** Relying on the results of Equation (4), the elastic net solution for the *j*^th^ loading vector is

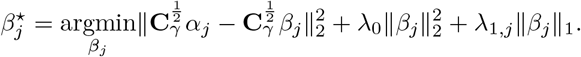

Generally, for ease of computation, λ_1,*j*_ = λ_1_, for *j* = 1,…, *k*. The entries of the loading matrix **B** are independent of the choice for the *ℓ*_2_ penalty (ridge) parameter λ_0_ [41], which is used only for numerical reasons. The ridge penalty is set to zero when 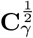 is of full rank; otherwise, a small constant value is used to remedy issues of indeterminacy that arise when fitting the elastic net.

**For fixed B:** Only the first term of the SPCA criterion of Equation (3) must be minimized with respect to **A**. The solution is given by the reduced rank form of the Procrustes rotation, computed as **A**⋆ = **UV**^⊤^ [41]. The matrices of left and right singular vectors are obtained from the following singular value decomposition:

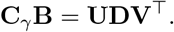

Generally, **C**_*γ*_ is not positive-semidefinite and its square root is undefined. Instead, a positive-semidefinite matrix 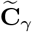, approximating **C**_*γ*_, is used. 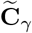 is obtained by replacing the diagonal matrix in the eigendecomposition of **C**_*γ*_ by a diagonal matrix in which negative eigenvalues are replaced by zeros [39]:

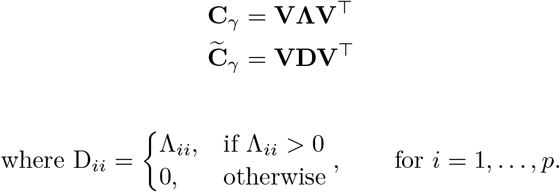

Thus, the directions of variation given by the negative eigenvalues of **C**_*γ*_ are discarded, as they correspond to those which are dominated by the variance in the background dataset. This procedure can be viewed as a preliminary thresholding of the eigenvectors of **C**_*γ*_, where the cutoff is an additional hyperparameter corresponding to a non-negative real number. Explicitly defining a small positive threshold may prove useful for datasets that possess many eigenvalues near zero, which correspond to sources of technical and biological noise remaining after the contrastive step. Empirically, however, providing a wide range of contrastive parameters *γ* has been found to have a similar effect as using multiple cutoff values, that is, larger values of *γ* naturally produce sparser matrices 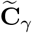.

For the purpose of contrastive analysis, a direction’s importance is characterized by its target-background variance coupling; higher target variance and lower background variance pairs produce the best directions [2] and correspond to the largest positive eigenvalues. The elimination of directions with negative eigenvalues therefore guarantees that the sparse contrastive PCs (scPCs) are rotations of the target data relying on the sparse directions most variable in the target data but least variable in the background data, making a cutoff of zero a natural choice for the thresholding operation.

### 2.2 Framework for Hyperparameter Tuning

The scPCA algorithm relies on two hyperparameters: the contrastive parameter *γ* and the *ℓ*_1_ penalty parameter λ_1_. To select the optimal combination of *γ* and λ_1_ from a grid of *a priori* specified values, we propose to cluster the *n* observations of the target dataset based on their first *k* scPCs, selecting as optimal the combination {*γ*, λ_1_} producing the “strongest” cluster assignments. This framework casts the selection of {*γ*, λ_1_} in terms of a choice of clustering algorithm, distance metric (based on 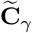), and clustering strength criterion. For ease of application, we propose to select {*γ*, λ_1_} by maximization of the average silhouette width over clusterings of the reduced-dimension representation of the target data. This procedure implicitly requires the choice of a clustering algorithm, such as *k*-means [20], to be applied to the representation of the data in the first *k* scPCs. Such methods require an appropriate choice for the number of clusters, which we contend will generally not be a limiting factor in the use of scPCA. Indeed, reasonable choices for the number of clusters can often be inferred in *omics* settings from sample annotation variables accompanying the data or from previously available biological knowledge. In Section 3, we empirically demonstrate that the results of the algorithm are robust to the choice of the number of clusters. Additionally, scPCA has no particular dependence on average silhouette width as a criterion, that is, alternative criteria for assessing clustering strength could be used when appropriate. Naturally, this proposed hyperparameter tuning approach can be applied to cPCA by setting λ_1_ to zero.

To address concerns of overfitting and to avoid discovering non-generalizable patterns from the data, we propose the use of cross-validation. For a grid of *a priori* specified contrastive parameters *γ* and *ℓ*_1_ penalty parameters λ_1_, *W*-fold cross-validation may be performed as follows:

1. Partition each of the target and background datasets into *W* roughly equally-sized subsets.
2. Randomly pair each of the target *W* subsets with one of the background subsets; these pairs form the fold-specific validation sets.
3. Iteratively perform scPCA over the observations of the target and background data not contained in the validation set (i.e., the training sets), for each pair of contrastive parameters and *ℓ*_1_ penalty parameters in the hyperparameter grid.
4. Project the target validation data onto the low-dimensional space using the loading matrices obtained in the prior step.
5. Compute a clustering strength criterion (e.g., average silhouette width) for a clustering of the target validation data with the *a priori* specified number of clusters.
6. Finally, compute the cross-validated average of the clustering strength criteria (e.g., cross-validated average of average silhouette width) across the validation sets for each pair of hyperparameters, selecting the pair that maximizes the value of the criterion.

We note two caveats of this cross-validation framework: (1) it increases the computation time, and (2) each fold is assumed to be representative of the process which generated the data. For large datasets and say 5 to 10 folds, (2) is not a grave concern; however, as the number of samples decreases, this assumption becomes less tenable. In such a situation, the non-cross-validated framework should be used, or the number of folds reduced. Examples and results are provided in Section S8.

### 2.3 Algorithm and Software Implementation

The implementation of the scPCA algorithm is presented in Algorithm 1 of the supplement. Algorithm 2, also in the supplement, details the cross-validated variant.

Regarding the computational complexity of scPCA (and cPCA), we note that the number of observations in the target and background data are not limiting factors. Indeed, these algorithms are applied to a *p* × *p* contrastive covariance matrix, a function of the target and background covariance matrices. The computation time of these covariance matrices increases linearly in the number of observations *n* and quadratically in the number of features *p*. The methods’ computational efficiency are therefore most impacted by the number of features and the size of the hyperparameter grid. A note on cPCA’s and scPCA’s running time is provided in Section S9, as is a comparison to competing methods.

A free and open-source software implementation of scPCA is available in the `scPCA` package for the `R` language and environment for statistical computing [26]. The `scPCA` package has been released as part of the Bioconductor Project [12, 11, 14] (https://bioconductor.org/packages/scPCA).

The code and data used to generate this manuscript are publicly available on GitHub (https://github.com/PhilBoileau/EHDBDscPCA).

## 3 Results

In the sequel, we detail the application of scPCA to a number of simulated and publicly available datasets, comparing our proposal to several competing techniques. An additional analysis of protein expression data is presented in Section S7. Details on hyperparameters used by all dimensionality reduction methods are presented in Sections S4, S5, and S6.

### 3.1 Simulated scRNA-seq Data

The scPCA technique was tested on a simulated scRNA-seq dataset generated with the *Splat* framework from the `Splatter R` package [36]. *Splat* simulates a scRNA-seq count matrix by way of a gamma-Poisson hierarchical model. This simulation framework mimics real scRNA-seq data by including hyperparameters to control the number of over- and under-expressed genes (using multiplicative factors for mean expression levels), zero inflation, batch effects, and other technical and biological factors relevant for scRNA-seq data.

A simple dataset of 300 cells and 500 genes was simulated such that the cells were approximately evenly distributed among three biological groups: two groups making up a target dataset and a third group corresponding to a background dataset. 5% of the genes are differentially expressed between the background dataset and each of the two target datasets but not between the two target datasets, 10% of the genes are differentially expressed between the background dataset and the first target dataset but not the second target dataset, and 10% of the genes are differentially expressed between the background dataset and the second target dataset but not the first target dataset. There is overlap between these three sets of genes and, in particular, a total of 98 genes are differentially expressed between the two target datasets. Based on these levels of differential expression, cells are more dissimilar between the two target datasets than between either of the target datasets and the background dataset. Therefore, the samples comprising the background dataset can be viewed as a set of controls for use by cPCA and scPCA. Additionally, a large batch effect was included to confound the biological variation between groups, effectively dividing each biological group into two subgroups of near equal size (Figure 1A).

**Figure 1:**
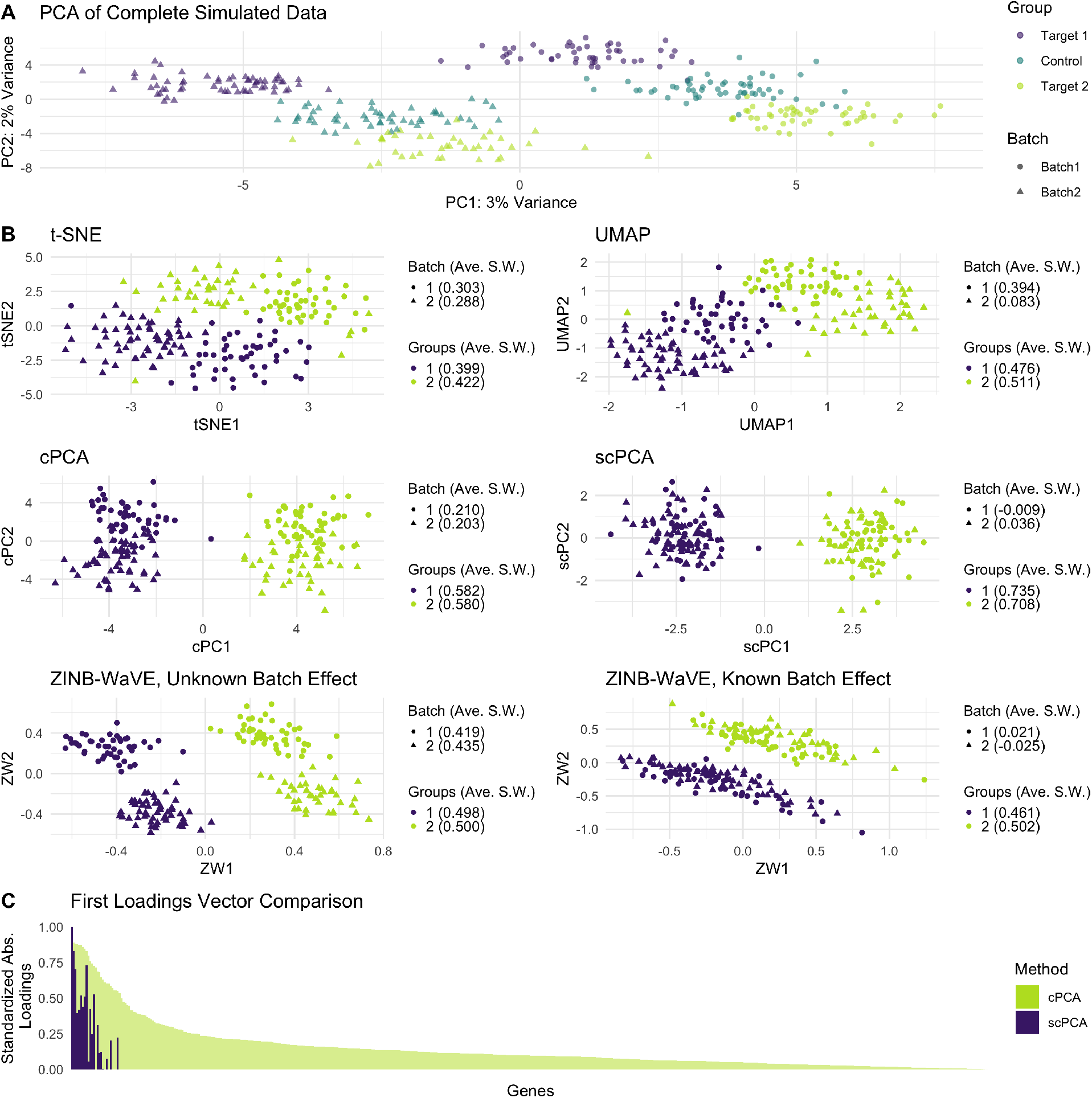
Simulated scRNA-seq data. **A** Plot of the first two principal components of the complete simulated dataset (i.e., the combination of the target and background datasets). The batch effect and the biological signal are responsible for approximately identical amounts of variance. **B** Two-dimensional representations of the target dataset by t-SNE, UMAP, cPCA, scPCA, and ZINB-WaVE, with accompanying average silhouette widths quantifying the strengths of the batch effect and the biological signal. Only scPCA fully removes the batch effect in two dimensions when batches are not adjusted for explicitly. **C** A gene-by-gene comparison of the standardized absolute loadings in the first loading vectors of cPCA and scPCA, in decreasing order with respect to the values produced by cPCA.

PCA, t-SNE, UMAP, cPCA, and scPCA were applied to the log-transformed and column-centered target data, and SIMLR to the raw target data, to determine whether these methods could identify the biological signal of interest, i.e., the two groups in the target dataset (Figure 1B. Note that PCA was not included due to the similarity of results to Figure 1A and SIMLR’s embedding is included in the appendix in Figure S4). Also note that cPCA was not performed in the traditional manner of Abid *et al*. [2], but with automatic hyperparameter selection as described in Sections 2.2 and 1. The number of *a priori* specified clusters for the cPCA and scPCA methods was set to 2, and the column-centered background data were used in their contrastive steps. While PCA, t-SNE, UMAP, SIMLR, and cPCA were incapable of completely eliminating the batch effect in their two-dimensional representations, scPCA successfully removed the unwanted variation while producing the tightest clusters, as indicated by the average silhouette widths (see also Figure S3), and generating sparse, interpretable loadings.

To compare the loadings produced by cPCA and scPCA, each of their loading vectors were standardized as follows. The *i*^th^ entry of the *j*^th^ standardized loading vector is given by 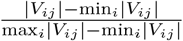, where **V** is a *p* × *k* loading matrix. Juxtaposing the standardized absolute loadings of the first loading vectors produced by cPCA and scPCA, each of which linearly separate the target dataset’s groups, we find the scPCA loadings to be, as expected, much sparser (see Figure 1C). In fact, only 20 genes have non-zero values in scPCA’s first loading vector compared to 500 in cPCA; moreover, these 20 genes correspond to those which have the largest absolute entries in cPCA’s first loading vector. Furthermore, these genes are among the most differentially expressed in the target dataset, based on the values of their multiplicative differential expression factors (Figure S2).

scPCA’s results were also compared to the two leading latent factors found by ZINB-WaVE, a method of choice for dimensionality reduction for scRNA-seq data, under conditions in which the batch factor is viewed as known and unknown (Figure 1B). In both cases, ZINB-WaVE was applied to the count matrix of the simulated target dataset with no gene-level covariates. When the batch factor was treated as unknown, no cell-level covariates were included in the model; however, when we treated the batch factor as known, a binary cell-level covariate was added to indicate each sample’s batch membership. When the batch effect is not explicitly regressed out in the ZINB-WaVE model, we find the results to be virtually identical to those of PCA. Even when the batch effect is included in the model, the clusters of the biological groups are elongated and less dense than those produced by scPCA, and the first latent factor does not linearly separate the groups.

### 3.2 Dengue Microarray Data

Kwissa *et al*. [18] used gene expression microarrays to analyze the whole-blood transcriptome of 47 dengue patients hospitalized at the Siriraj Hospital in Bangkok and 9 local healthy controls. Of the affected patients, 18 were classified as having acute dengue fever (DF), 10 as having acute dengue hemorrhagic fever (DHF), and 19 as convalescent at least four weeks after discharge.

As part of data pre-processing, all but the 500 most variable genes were filtered out. The target dataset consists of the log-transformed microarray expression measures of 47 patients with some form of dengue, while the background dataset consist of the log-transformed microarray expression measures of the control samples. PCA, cPCA, scPCA, t-SNE, and UMAP were then applied to the column-centered target data matrix with the goal of discerning three unique clusters (Figure 2A), one for each sub-class of dengue (DF, DHF, and convalescent). cPCA and scPCA took as additional input the column-centered background data matrix and specified three clusters *a priori*. t-SNE’s embedding was found to be similar to UMAP’s and is therefore only included in the supplementary materials (Figure S5).

**Figure 2:**
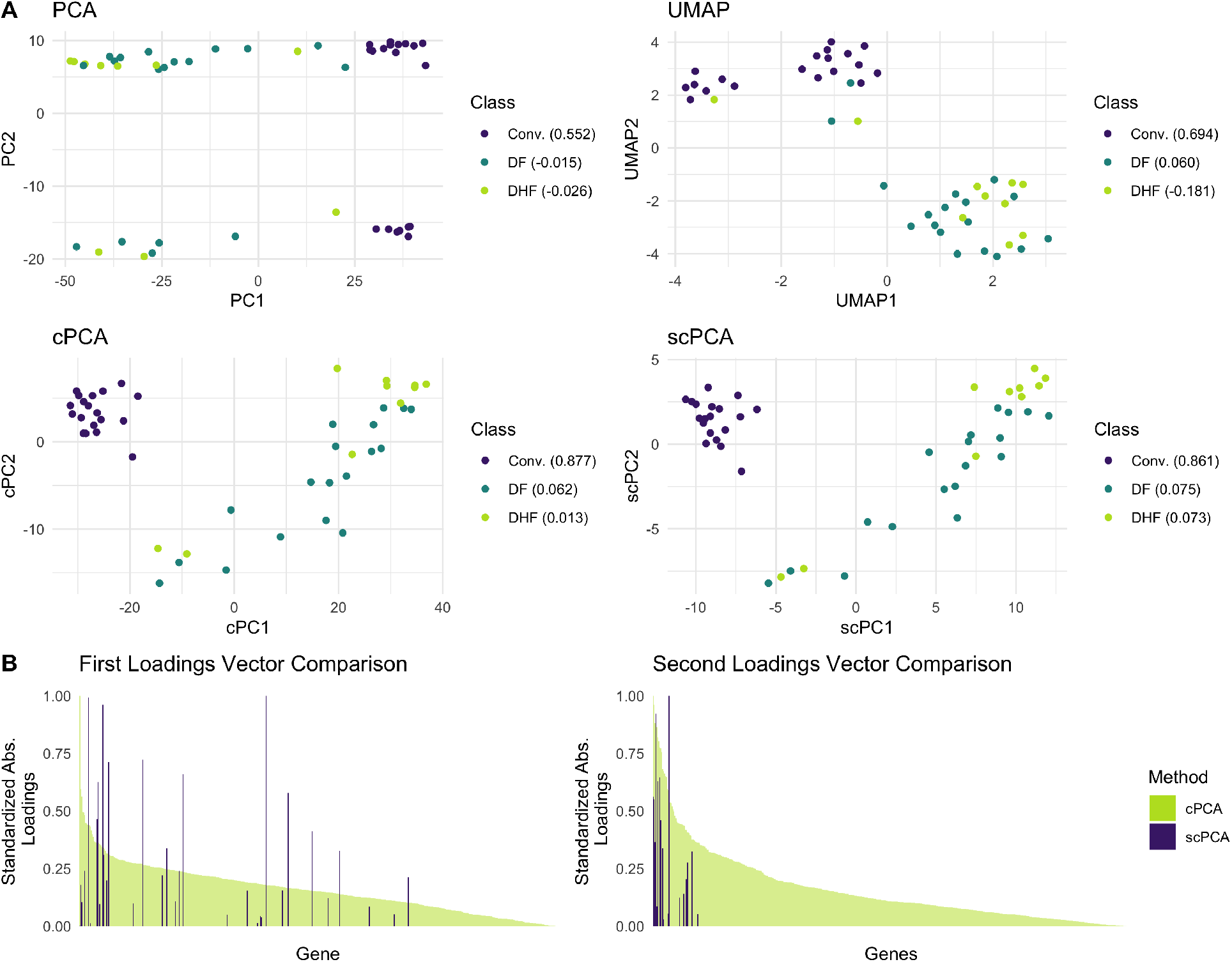
Dengue microarray data. **A** Two-dimensional representations of the target dataset by PCA, UMAP, cPCA, and scPCA, with accompanying average silhouette widths quantifying the strengths of the biological signal. cPCA and scPCA are the only methods that fully separate the convalescent patients from those with DF and DHF. The second PC of the PCA plot is dominated by some batch effect, and the low-dimensional representation produced by UMAP also appears to be affected by some source of unwanted variation. **B** The standardized absolute loadings in the two leading loading vectors of scPCA are much sparser than those of cPCA, though their two-dimensional embeddings are virtually identical. The genes are in decreasing order of cPCA’s standardized absolute loadings, demonstrating that the genes with non-zero loadings in scPCA generally correspond to the genes with the largest absolute loadings in cPCA. This is much more apparent for the second loading vector where the distribution of cPCA’s absolute loadings has a thin tail, attributing increased importance to a small subset of genes.

Of the four dimensionality reduction methods, only cPCA and scPCA successfully fully separated the convalescent patients from those with DF and DHF in two dimensions. scPCA’s low-dimensional representation was virtually identical to that of cPCA, producing very similar average silhouette widths among classes, though only a tenth of the genes have non-zero values in the first and second columns of the scPCA loading matrix, and the most important genes identified by each methods’ first loading vector differ substantially (Figure 2B). The genes found by scPCA include CD38, HLA-DQB1, and RSAD2 (Viperin), which have been previously associated to the susceptibility to, protection against, or presence of dengue [6, 5, 7]. For a full list of these genes, refer to Table S1 and Table S2. A gene set enrichment analysis (GSEA) was also performed with the genes contained in these tables to identify the most significant biological processes in which they play a role; details and results are presented in Table S3.

No method successfully distinguished between the three sub-classes of dengue. In fact, previous research suggests that the transcriptomes of patients with DF and DHF are virtually indistinct [18]. Instead, Kwissa *et al*. [18] found that DF and DHF patients may form distinct clusters based on viral load and concentration of the DENV NS-1 antigen in their plasma. This may explain the sub-clusters within the DF and DHF cases found by UMAP. Though the number of pre-specified clusters for each algorithm was set to three, cPCA’s and scPCA’s projections onto two dimensions contain two clusters. To test the sensitivity of these methods to this tuning parameter, both methods were reapplied to the data with varying numbers of pre-specified clusters. Each of cPCA’s iteration produced virtually identical embeddings (Figure S6). However, scPCA’s produced identical results to those of PCA when the number of clusters was set to four or higher (Figure S7). This may provide an empirical approach to selecting the appropriate number of clusters for scPCA, i.e., selecting the largest value before which the quality of the embedding deteriorates.

### 3.3 Leukemia Patient scRNA-seq Data

Finally, we tested scPCA on scRNA-seq data from the cryopreserved bone marrow mononuclear cell (BMMC) samples of two acute myeloid leukemia (AML) patients (Patient 035: 4,501 cells; Patient 027: 7,898 cells), before and after undergoing allogeneic hematopoietic stem cell transplant treatment [38]. The BMMCs of two healthy individuals from the same publicly available dataset (Healthy 1: 1,985 cells; Healthy 2: 2,472 cells) were used to generate a control dataset. Following pre-processing, all but the 1,000 most variable genes measured across all 16,856 cells were removed. The scRNA-seq data from the AML patients were then split into separate target datasets since Zheng *et al*. [38] found evidence of distinct subpopulation membership following transplantation. Data belonging to the healthy controls were combined to create the background dataset. PCA, t-SNE, UMAP, ZINB-WaVE, SIMLR, cPCA, and scPCA were applied to the target datasets to explore differences in the AML patients’ BMMCs’s transcriptome engendered by the treatment (Figures 3, S9, S8, and S10).

**Figure 3:**
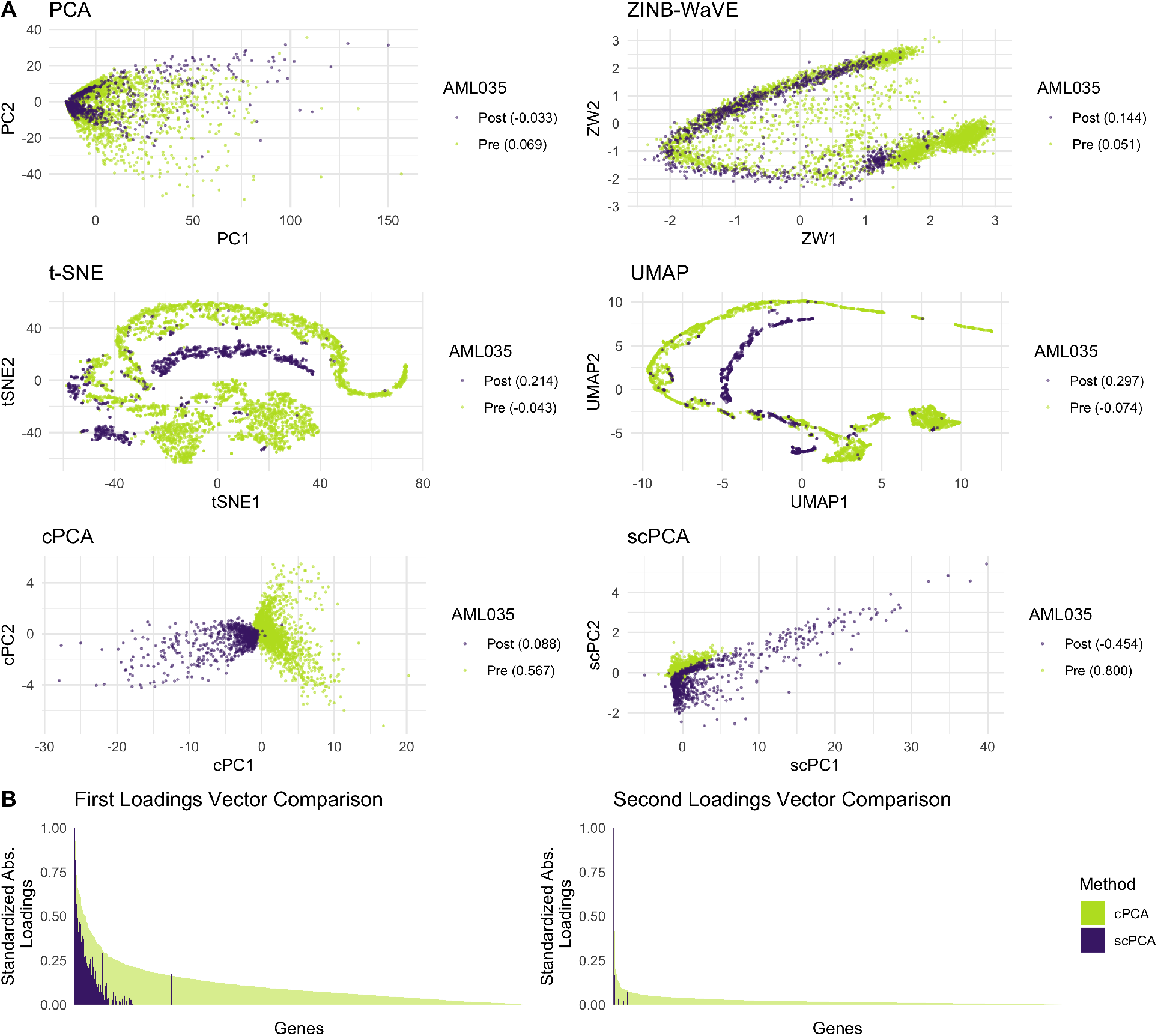
AML Patient 035 scRNA-seq data. **A** The two-dimensional embeddings of the patient’s BMMCs produced by PCA, ZINB-WaVE, t-SNE, UMAP, cPCA, and scPCA, with accompanying average silhouette widths quantifying the strengths of the biological signal. cPCA and scPCA produce representations of the data in which the pre- and post-transplant cells form discernible clusters. Based on visual inspection and average silhouette width, scPCA’s grouping of pre-transplant cells is denser than that of cPCA’s and the opposite is true of the post-transplant cells’ cluster. **B** scPCA’s embedding is much sparser, increasing interpretability of the exploratory analysis.

Of the seven dimensionality reduction methods applied to Patient 035’s data (Figures 3A and S8), cPCA and scPCA best capture the biologically meaningful information relating to treatment status. Each produces linearly separable clusters corresponding to pre- and post-treatment cells; scPCA’s projection yields a tighter cluster of pre-transplant cells when compared to that produced by cPCA, and the opposite is true regarding the clusters of post-transplant cells. Additionally, scPCA’s projection required considerably less information, even though its results are analogous to cPCA’s: 176 genes and 17 genes have non-zero entries in, respectively, the first and second columns of the loading matrix produced by scPCA (Figure 3B). In general, the leading loading vectors of cPCA and scPCA place an increased importance on the same genes. Genes with non-zero loadings in scPCA’s first and second loading vectors were also subjected to a gene set enrichment analysis to uncover their roles in biological processes; details and results are presented in Table S4. Regarding the other methods’ results, PCA, t-SNE, and SIMLR fail to separate the pre- and post-transplant cells, and UMAP’s and ZINB-WaVE’s embeddings resemble a trajectory more closely than they do a set of clusters.

Similarly to Patient 035’s results, the two-dimensional embeddings of Patient 027’s data produced by PCA, t-SNE, UMAP, SIMLR, and ZINB-WaVE do not contain distinct clusters of pre- and post-transplant BMMCs S9; however, cPCA and scPCA generate low-dimensional representations of the data in which samples are clustered based on treatment status. Although cPCA’s representation produces denser, more distinct groupings, the first two columns of scPCA’s loading matrix contain non-zero values in only five genes, STMN1, CA1, LDHA, PDLIM1, and C1QBP. The first four genes have been linked to leukemia [22, 24, 34, 13], and the last gene, C1QBP, is responsible for a protein that plays a critical role in tumor metabolism [1].

## 4 Discussion

We have proposed a novel dimensionality reduction technique for use with high-dimensional biological data: *sparse contrastive principal component analysis*. A central contribution of the method is the incorporation of a penalization step that ensures both the sparsity and stability of the principal components generated by contrastive PCA. Our approach allows for high-dimensional biological datasets, such as those produced by the currently popular scRNA-seq experimental paradigm, to be examined in a manner that uncovers interpretable as well as relevant biological signal after the removal of unwanted technical variation, all the while placing only minimal assumptions on the data-generating process.

We also present a data-adaptive and algorithmic framework for applying contrastive dimensionality reduction techniques, like cPCA and scPCA, to high-dimensional biological data. Where the original proposal of cPCA relied upon visual inspection by the user in selecting the “best” contrastive parameter [2], an unreliable process, our extension formalizes the data-adaptive selection of tuning parameters. The automation of this step translates directly to significantly increased computational reproducibility. We have proposed the use of cross-validation to select tuning parameters from among a pre-specified set in a generalizable manner, using average silhouette width to assess clustering strength. Several other approaches to the selection of tuning parameters, including the choice of criterion for assessing the “goodness” of the dimensionality reduction (here, clustering strength as measured by the average silhouette width), may outperform our approach in practice and could be incorporated into the modular framework of scPCA; we leave the development of such approaches and assessment of their potential advantages as an avenue for future investigation.

We have demonstrated the utility of scPCA relative to competing approaches, including standard PCA, cPCA, t-SNE, UMAP, and, where appropriate, ZINB-WaVE and SIMLR, using both a simulation study of single-cell RNA-seq data and the re-analysis of several publicly available datasets from a variety of high-dimensional biological assays. We have shown that scPCA recovers low-dimensional embeddings similar to cPCA, but with a more easily interpretable principal component structure and, in simulations, diminished technical noise. Further, our results indicate that scPCA generally produces denser, more relevant clusters than t-SNE, UMAP, SIMLR, and ZINB-WaVE. Moreover, we verify that clusters derived from scPCA correspond to biological signal of interest. Finally, as the cost of producing high-dimensional biological data with high-throughput experiments continues to decrease, we expect that the availability and utility of techniques like scPCA – for reliably extracting rich, sparse biological signals while data-adaptively removing technical artifacts – will strongly motivate the collection of control samples as a part of standard practice.

## Acknowledgements

We thank Mark van der Laan and Hector Roux de Bézieux for helpful discussions and insights.

