## Supplementary Information for "Exploring High-Dimensional Biological Data with Sparse Contrastive Principal Component Analysis"

A Preprint

**Philippe Boileau**
Graduate Group in Biostatistics,
University of California, Berkeley


**Nima S. Hejazi**
Graduate Group in Biostatistics and
Center for Computational Biology,
University of California, Berkeley


**Sandrine Dudoit**
Department of Statistics,
Division of Biostatistics, and
Center for Computational Biology,
University of California, Berkeley


February 22, 2020

### S1 Algorithm for scPCA

**Algorithm 1**: scPCA

**Result**: Produces a sparse low-dimensional representation of the target data, $\mathbf{X}_{n \times p}$, by contrasting the variation of $\mathbf{X}_{n \times p}$ and some background data, $\mathbf{Y}_{m \times p}$, while applying an $\ell_1$ penalty to the loadings generated by cPCA.

**Input** :

target dataset: $\mathbf{X}$

background dataset: $\mathbf{Y}$

binary variable indicating whether to column-scale the data: `scale`

vector of possible contrastive parameters: $\gamma = (\gamma_1, \ldots, \gamma_s)$

vector of possible $\ell_1$ penalty parameters: $\lambda_1 = (\lambda_{1,1}, \ldots, \lambda_{1,d})$

number of sparse contrastive principal components to compute: $k$

clustering method: `cluster_meth`

number of clusters: `ncluster`

Center (and `scale` if so desired) the columns of $\mathbf{X}$, $\mathbf{Y}$

Calculate the empirical covariance matrices: $\mathbf{C}_{\mathbf{X}_{p \times p}} := \frac{1}{n}\mathbf{X}^\top\mathbf{X}$, $\mathbf{C}_{\mathbf{Y}_{p \times p}} := \frac{1}{m}\mathbf{Y}^\top\mathbf{Y}$

**for** *each* $\gamma_i \in \gamma$ **do**

  **for** *each* $\lambda_{1,j} \in \lambda_1$ **do**

    Compute the contrastive covariance matrix $\mathbf{C}_{\gamma_i} = \mathbf{C}_\mathbf{X} - \gamma_i \mathbf{C}_\mathbf{Y}$

    Compute the positive-semidefinite approximation of $\mathbf{C}_{\gamma_i}$, $\widetilde{\mathbf{C}}_{\gamma_i}$

    Apply SPCA to $\widetilde{\mathbf{C}}_{\gamma_i}$ for $k$ components with $\ell_1$ penalty $\lambda_{1,j}$

    Generate a low-dimensional representation by projecting $\mathbf{X}_{n \times p}$ on the sparse loadings of SPCA

    Normalize the low-dimensional representation produced to be on the unit hypercube

    Cluster the normalized low-dimensional representation using `cluster_meth` with `ncluster`

    Compute and record the clustering strength criterion associated with $(\gamma_i, \lambda_{1,j})$

Identify the combination of hyperparameters maximizing the clustering strength criterion: $\gamma^\star$, $\lambda_1^\star$

**Output**: The low-dimensional representation of the target data given by $(\gamma^\star, \lambda_1^\star)$, an $n \times k$ matrix; the $p \times k$ matrix of loadings given by $(\gamma^\star, \lambda_1^\star)$; contrastive parameter $\gamma^\star$; $\ell_1$ penalty parameter $\lambda_1^\star$

### S2 Algorithm for Cross-Validated scPCA

**Algorithm 2**: Cross-validated scPCA

**Result**: Produces a sparse low-dimensional representation of the target data, $\mathbf{X}_{n \times p}$, by contrasting the variation of $\mathbf{X}_{n \times p}$ and some background data, $\mathbf{Y}_{m \times p}$, while applying an $\ell_1$ penalty to the loadings generated by cPCA.

**Input** :

target dataset: $\mathbf{X}$

background dataset: $\mathbf{Y}$

binary variable indicating whether to column-scale the data: `scale`

vector of possible contrastive parameters: $\gamma = (\gamma_1, \ldots, \gamma_s)$

vector of possible $\ell_1$ penalty parameters: $\lambda_1 = (\lambda_{1,1}, \ldots, \lambda_{1,d})$

number of sparse contrastive principal components to compute: $k$

clustering method: `cluster_meth`

number of clusters: `ncluster`

number of cross-validation folds: $W$

For $\mathbf{X}_{n \times p}$, randomly partition the index set $\{1, \ldots, n\}$ into $W$ validation sets, $\mathcal{W}_1^x, \ldots, \mathcal{W}_W^x$, of (approximately) the same size (i.e., $\bigcup_{w=1}^{W} \mathcal{W}_w^x = \{1, \ldots, n\}$; $\mathcal{W}_w^x \cap \mathcal{W}_{w'}^x = \emptyset, \quad \forall w, w' \in \{1, \ldots, W\}$). Denote the corresponding training sets by $\mathcal{T}_w^x = \{1, \ldots, n\} \setminus \mathcal{W}_w^x$. For $\mathbf{Y}_{m \times p}$, randomly partition the index set $\{1, \ldots, m\}$ into $W$ validation sets, $\mathcal{W}_1^y, \ldots, \mathcal{W}_W^y$, of (approximately) the same size (i.e., $\bigcup_{w=1}^{W} \mathcal{W}_w^y = \{1, \ldots, m\}$; $\mathcal{W}_w^y \cap \mathcal{W}_{w'}^y = \emptyset, \quad \forall w, w' \in \{1, \ldots, W\}$). Denote the corresponding training sets by $\mathcal{T}_w^y = \{1, \ldots, m\} \setminus \mathcal{W}_w^y$. Denote by $\mathbf{X}_{\mathcal{T}_w^x}$ the $|\mathcal{T}_w^x| \times p$ submatrix of $\mathbf{X}$ for training set $\mathcal{T}_w^x$ and by $\mathbf{Y}_{\mathcal{T}_w^y}$ the $|\mathcal{T}_w^y| \times p$ submatrix of $\mathbf{Y}$ for training set $\mathcal{T}_w^y$. Define similarly $\mathbf{X}_{\mathcal{W}_w^x}$ and $\mathbf{Y}_{\mathcal{W}_w^y}$ for the validation sets. Note that $\mathbf{Y}_{\mathcal{W}_w^y}$ is defined explicitly solely to avoid ambiguity; it plays no role in subsequent developments.

**for** *each* $w$ *in* $\{1, \ldots, W\}$ **do**
- Center (and `scale` if so desired) the columns of $\{\mathbf{X}_{\mathcal{T}_w^x}, \mathbf{Y}_{\mathcal{T}_w^y}\}$ and $\{\mathbf{X}_{\mathcal{W}_w^x}, \mathbf{Y}_{\mathcal{W}_w^y}\}$
- Compute the empirical covariance matrices: $\mathbf{C}_{\mathbf{X}_{p \times p}} := \frac{1}{|\mathcal{T}_w^x|} \mathbf{X}_{\mathcal{T}_\mathbf{w}^\mathbf{x}}{}^\top \mathbf{X}_{\mathcal{T}_\mathbf{w}^\mathbf{x}}$, $\mathbf{C}_{\mathbf{Y}_{p \times p}} := \frac{1}{|\mathcal{T}_w^y|} \mathbf{Y}_{\mathcal{T}_\mathbf{w}^\mathbf{y}}{}^\top \mathbf{Y}_{\mathcal{T}_\mathbf{w}^\mathbf{y}}$
- **for** *each* $\gamma_i \in \gamma$ **do**
  - **for** *each* $\lambda_{1,j} \in \lambda_1$ **do**
    - Compute the contrastive covariance matrix $\mathbf{C}_{\gamma_i} = \mathbf{C}_\mathbf{X} - \gamma_i \mathbf{C}_\mathbf{Y}$
    - Compute the positive-semidefinite approximation of $\mathbf{C}_{\gamma_i}$, $\widetilde{\mathbf{C}}_{\gamma_i}$
    - Apply SPCA to $\widetilde{\mathbf{C}}_{\gamma_i}$ for $k$ components with $\ell_1$ penalty $\lambda_{1,j}$
    - Generate a low-dimensional representation of the target validation set by projecting $\mathbf{X}_{\mathcal{W}_w^x}$ on the sparse loadings of SPCA
    - Normalize the low-dimensional representation produced to be on the unit hypercube
    - Cluster the normalized low-dimensional representation using `cluster_meth` with `ncluster`
    - Compute and record the clustering strength criterion associated with $(\gamma_i, \lambda_{1,j})$

Identify the combination of hyperparameters maximizing the cross-validated mean (across all folds $\{1, \ldots, W\}$) of the clustering strength criterion: $\gamma^\star$, $\lambda_1^\star$

**Output**: The low-dimensional representation of the target data given by $(\gamma^\star, \lambda_1^\star)$, a $n \times k$ matrix; the $p \times k$ matrix of loadings given by $(\gamma^\star, \lambda_1^\star)$; contrastive parameter $\gamma^\star$; $\ell_1$ penalty parameter $\lambda_1^\star$

#### S3 Intuition for the Contrastive Parameter $\gamma$

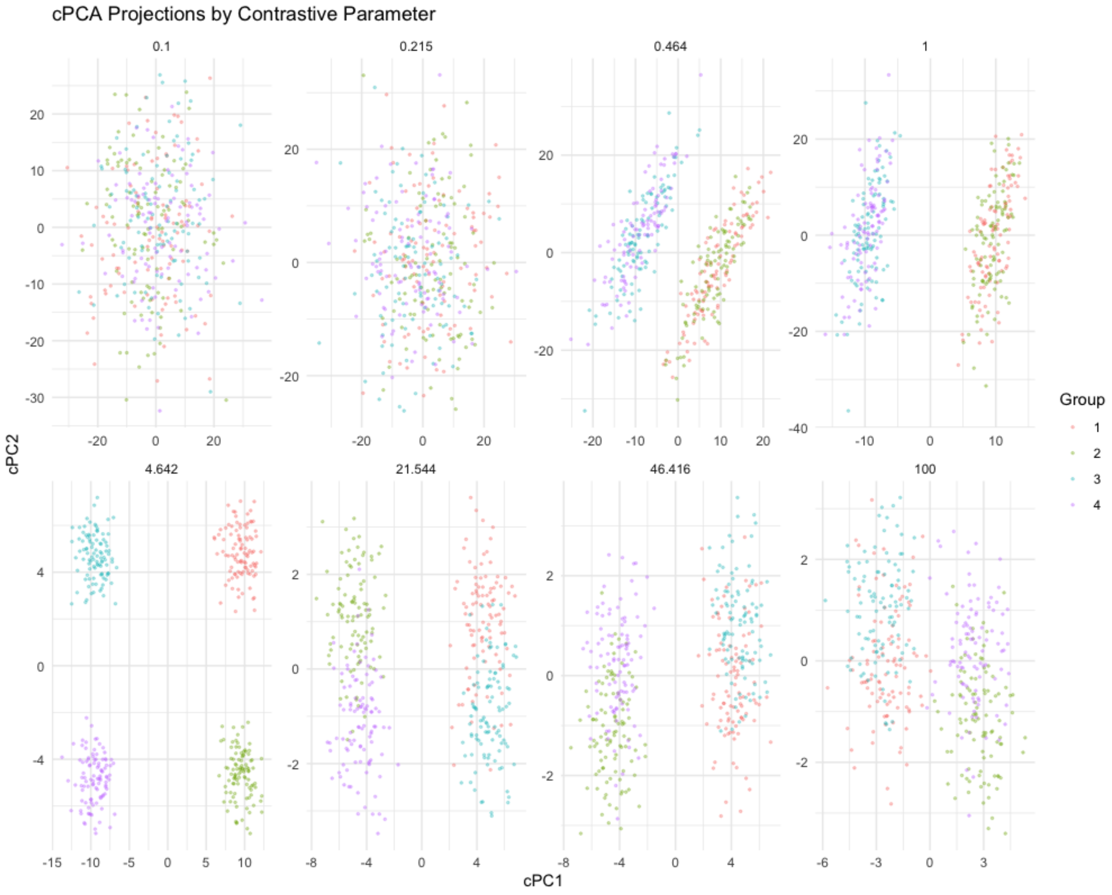


Figure S1: *Effect of contrastive parameter for cPCA.* cPCA as implemented by Abid *et al.* [1] was applied to a simulated dataset of $n = 400$ observations, split across 4 groups, with $p = 30$ variables. The first 10 variables are distributed as $N(0, 10)$ for all observations. Variables 11 through 20 are distributed as $N(0, 1)$ for Groups 1 an 2, and as $N(3, 1)$ for Groups 3 and 4. Variables 21 through 30 are distributed as $N(-3, 1)$ for Groups 1 and 3, and as $N(0, 1)$ for Groups 2 and 4. cPCA also takes as input a background dataset of $m = 400$ observations, with $p = 30$ variables, where the first 10 variables are distributed as $N(0, 10)$, the following 10 as $N(0, 3)$, and the remaining 10 as $N(0, 1)$. The results of cPCA are then presented for eight increasing values of the contrastive parameter $\gamma$ from among 40 loagrithmically spaced values between 0.1 and 100, selected using the semi-automated technique described by Abid *et al.* [1]. For the smaller values of the contrastive parameter, the noise contained in the first 10 variables of the target data dominates the signal contained in variables 11 through 30. As the contrastive parameter increases, the signal in the target data set is unmasked. However, once the contrastive parameter value becomes larger than $\approx 20$, the distinction between groups becomes increasingly poor; the variation contained in the background data begins to dominate the variation contained in the target data. A virtually identical dataset is presented in the supplementary material of Abid *et al.* [1].

#### S4 Simulated scRNA-seq Data

See Section 3.1 for information on the simulation model and dataset. cPCA was applied to the dataset using the non-cross-validated hyperparameter tuning framework. The $\ell_1$ penalty parameter was set to 0, and the vector of possible contrastive parameters consisted of 40 logarithmically spaced values between 0.1 and 1,000. scPCA was applied in the same manner as cPCA, though the vector of potential $\ell_1$ penalty parameters consisted of 20 equidistant values between 0.05 and 1. The `Rtsne R` package was used to create the t-SNE embedding. Two initializations were generated, that is, with and without an initial application of PCA to the simulated data. The embedding employing the initial PCA step retained the 50 leading principal components. For each initialization, the remaining parameters were set to their defaults, as done by Becht *et al.* [3], e.g., `perplexity = 30` and `max_iter = 1000`. The `theta` parameter was set to `0` so that exact t-SNE was performed. The embedding produced using the PCA initialization was qualitatively better than that without. Therefore, it is used in Figure 1 of this manuscript. The `umap R` package was used to generate the UMAP embedding. As in the quantitative analysis of UMAP performed by Becht *et al.* [3], `min_dist` was set to `0.02`, `nearest_neighbors` was set to `30`, and the Euclidean distance was used as a metric. The SIMLR was produced with a `k` of 10, as recommended in [15], and setting the number of pre-specified clusters to 2.

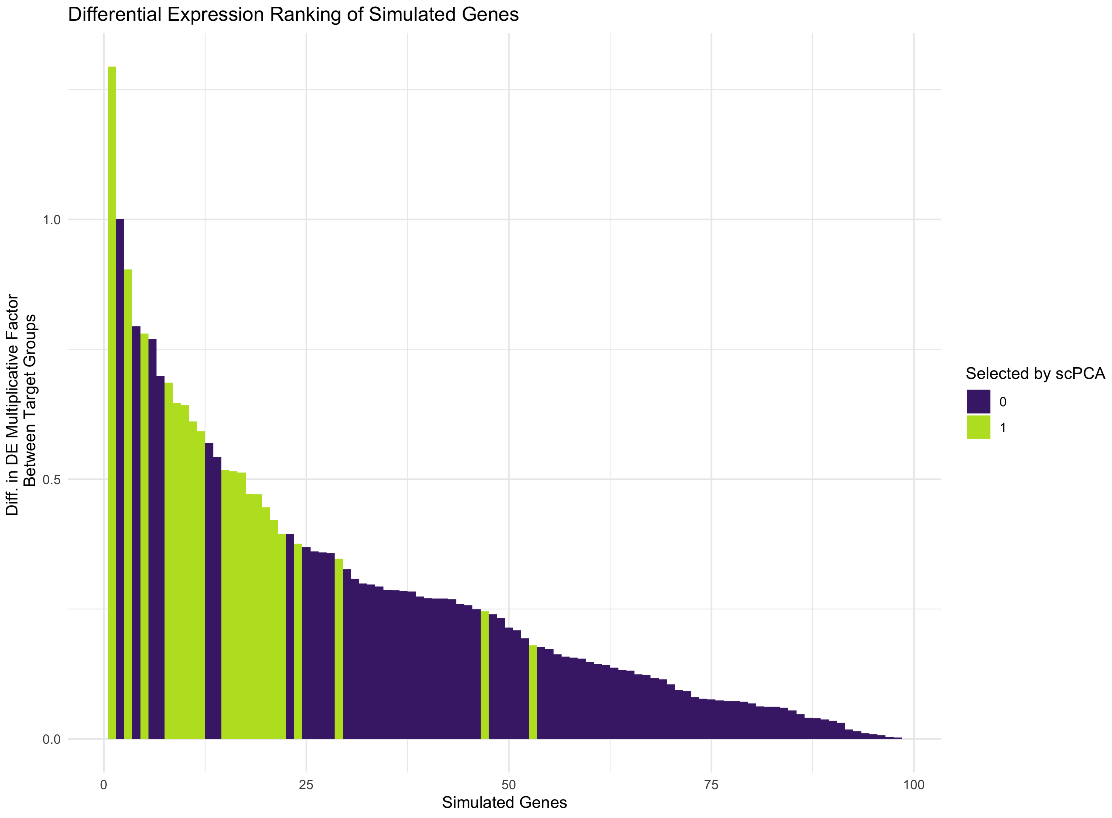


Figure S2: *Simulated scRNA-seq data: Differential expression.* The 98 differentially expressed genes in the simulated target dataset are ranked in decreasing order of their absolute level of differential expression between groups. In the *Splatter* framework, genes are differentially expressed between groups by way of a group-specific multiplicative factor. Thus, the level of differential expression of any gene between two groups may be computed as the absolute value of the difference between each group's multiplicative factor. We find that all 20 of the genes with non-zero entries in scPCA's first loading vector, highlighted in green, are among the most differentially expressed.

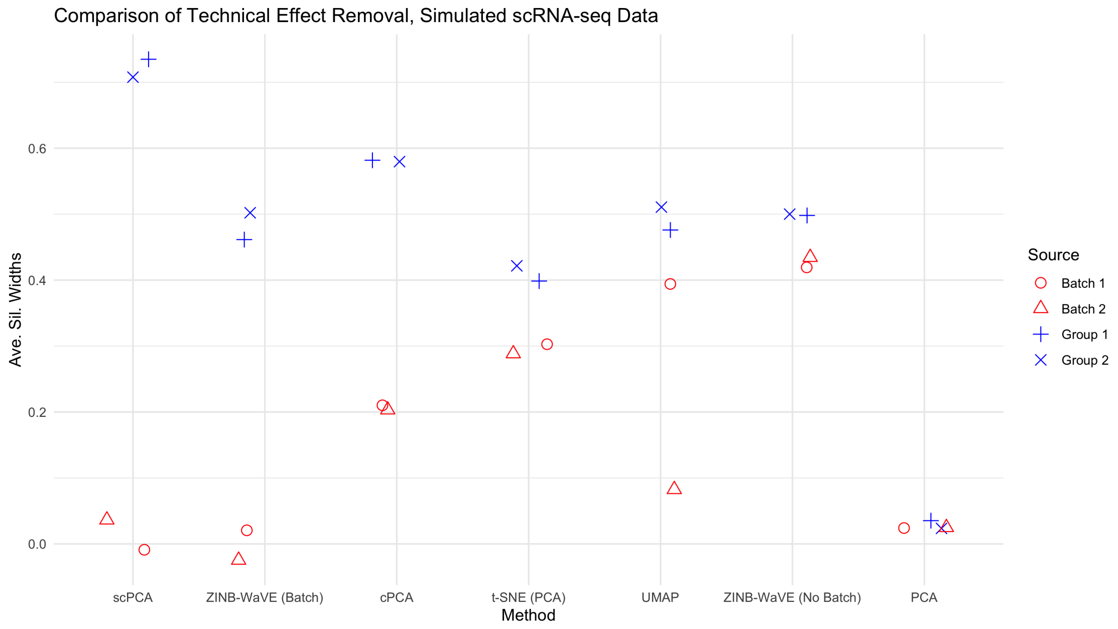


Figure S3: *Simulated scRNA-seq data: Average silhouette width comparison.* Methods are displayed in decreasing order of ability to remove unwanted technical variation, as measured by the average silhouette width. scPCA produces the densest biological clusters with the least amount of technical noise. The ZINB-WaVE method, when taking into account the batch effect, has a similar performance to scPCA with respect to the removal of unwanted effects, though the biological clusters it produces have lower average silhouette widths. Though cPCA produces denser biological clusters than ZINB-WaVE, it fails to completely remove the batch effect. The remaining methods are unable to disentangle the biological and technical effects.

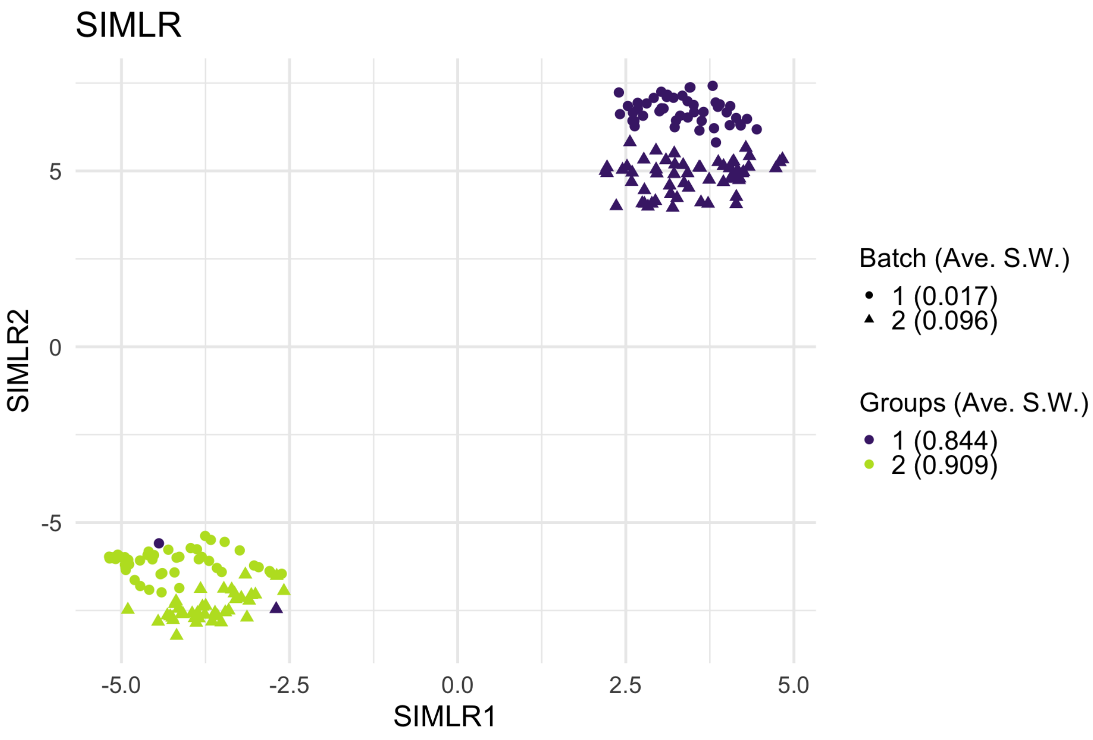


Figure S4: *Simulated scRNA-seq data: SIMLR.* SIMLR's two-dimensional embedding produces dense clusters that nearly perfectly split cells into two biologically meaningful groups. However, upon close inspection – and contrary to what the average silhouette widths suggest – the batch effect is not removed from the biological groups. Because the reported average silhouette widths misrepresent the method's ability to remove unwanted variation in the data, its results were not included in Figure S3.

#### S5 Dengue Microarray Data

See Section 3.2 for information on the data. cPCA and scPCA were fit using the non-cross-validated hyperparameter tuning framework. Both methods considered a vector of 40 logarithmically spaced values between 0.1 and 1,000 as potential contrastive parameters. scPCA also used a vector of 20 equidistant values between 0.05 and 1 as potential $\ell_1$ penalty parameters. As with the simulated data, two t-SNE embeddings were generated: one with an initial PCA step (retaining the first 50 principal components) and one without. Due to the small sample size, the `perplexity` parameter was set to 8. The remaining hyperparameters were set to their defaults, with the exception of `theta` which was set to 0. Both embeddings were qualitatively identical, and so only that which does not require the initial dimensionality reduction through PCA is presented in this manuscript. The qualitatively best UMAP embedding was found with the `n_neighbors` parameter set to 15 and the `min_dist` parameter set to 0.2. These parameter values are inspired from those used by Becht *et al.* [3], though the type of data considered are not identical.

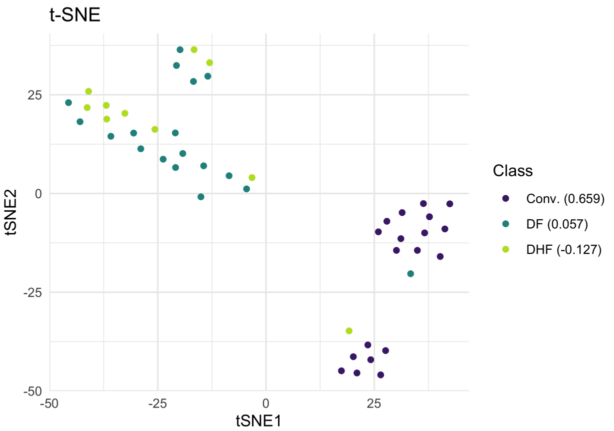


Figure S5: *Dengue microarray data: t-SNE.* Similarly to UMAP, t-SNE almost completely separates the convalescent patients from those with some form of dengue. The two main clusters are further split into distinct sub-clusters, perhaps indicating the presence of a batch effect.

Table S1: *Dengue microarray data: Genes with non-zero weights in the first scPCA loading vector.*

| | Gene Symbol | Gene Name | Weight |
|---|---|---|---|
| 1 | PRSS33 | protease, serine, 33 | -0.0059 |
| 2 | PDZK1IP1 | PDZK1 interacting protein 1 | -0.0347 |
| 3 | SDC1 | syndecan 1 | 0.2507 |
| 4 | CAV1 | caveolin 1, caveolae protein, 22kDa | 0.0889 |
| 5 | GGH | gamma-glutamyl hydrolase (conjugase, folylpolygammaglutamyl hydrolase) | 0.2318 |
| 6 | PI3 | peptidase inhibitor 3, skin-derived | -0.0209 |
| 7 | BUB1B | budding uninhibited by benzimidazoles 1 homolog beta (yeast) | 0.1242 |
| 8 | ZWINT | ZW10 interactor | 0.3984 |
| 9 | TUBB2A | tubulin, beta 2A | -0.0004 |
| 10 | PTGS2 | prostaglandin-endoperoxide synthase 2 (prostaglandin G/H synthase and cyclooxygenase) | -0.0627 |
| 11 | TTK | TTK protein kinase | 0.0201 |
| 12 | ORM1 /// ORM2 | orosomucoid 1 /// orosomucoid 2 | -0.0055 |
| 13 | CD38 | CD38 molecule | 0.0399 |
| 14 | CHI3L1 | chitinase 3-like 1 (cartilage glycoprotein-39) | -0.0384 |
| 15 | HLA-DQB1 | major histocompatibility complex, class II, DQ beta 1 | 0.0720 |

Table S1: *Dengue microarray data: Genes with non-zero weights in the first scPCA loading vector.*

| | Gene Symbol | Gene Name | Weight |
|---|---|---|---|
| 16 | BUB1 | budding uninhibited by benzimidazoles 1 homolog (yeast) | 0.0853 |
| 17 | CDK1 | cyclin-dependent kinase 1 | 0.2650 |
| 18 | IGH@ /// IGHA1 /// IGHA2 /// IGHD /// IGHG1 /// IGHG3 /// IGHG4 /// IGHM /// IGHV4-31 /// LOC100290146 /// LOC100290528 | immunoglobulin heavy locus /// immunoglobulin heavy constant alpha 1 /// immunoglobulin heavy constant alpha 2 (A2m marker) /// immunoglobulin heavy constant delta /// immunoglobulin heavy constant gamma 1 (G1m marker) /// immunoglobulin heavy constant gamma 3 (G3m marker) /// immunoglobulin heavy constant gamma 4 (G4m marker) /// immunoglobulin heavy constant mu /// immunoglobulin heavy variable 4-31 /// hypothetical protein LOC100290146 /// similar to pre-B lymphocyte gene 2 | 0.0180 |
| 19 | IGH@ /// IGHA1 /// IGHA2 /// IGHD /// IGHG1 /// IGHG3 /// IGHG4 /// IGHM /// IGHV3-23 /// LOC100126583 /// LOC100290146 /// LOC652128 | immunoglobulin heavy locus /// immunoglobulin heavy constant alpha 1 /// immunoglobulin heavy constant alpha 2 (A2m marker) /// immunoglobulin heavy constant delta /// immunoglobulin heavy constant gamma 1 (G1m marker) /// immunoglobulin heavy constant gamma 3 (G3m marker) /// immunoglobulin heavy constant gamma 4 (G4m marker) /// immunoglobulin heavy constant mu /// immunoglobulin heavy variable 3-23 /// hypothetical LOC100126583 /// hypothetical protein LOC100290146 /// similar to Ig heavy chain V-II region ARH-77 precursor | 0.1867 |
| 20 | NOV | nephroblastoma overexpressed gene | -0.0619 |
| 21 | SELENBP1 | selenium binding protein 1 | -0.1315 |
| 22 | IGHA1 /// IGHG1 /// IGHM /// LOC100290293 | immunoglobulin heavy constant alpha 1 /// immunoglobulin heavy constant gamma 1 (G1m marker) /// immunoglobulin heavy constant mu /// similar to hCG2042717 | 0.0162 |
| 23 | CEP55 | centrosomal protein 55kDa | 0.2863 |
| 24 | PBK | PDZ binding kinase | 0.1358 |
| 25 | SHCBP1 | SHC SH2-domain binding protein 1 | 0.2901 |
| 26 | MGC29506 | plasma cell-induced ER protein 1 | 0.4012 |
| 27 | CNTNAP3 | contactin associated protein-like 3 | -0.0494 |
| 28 | JAZF1 | JAZF zinc finger 1 | -0.0441 |
| 29 | KIAA1324 | KIAA1324 | -0.0962 |
| 30 | CDCA2 | cell division cycle associated 2 | 0.3858 |
| 31 | KLHL14 | kelch-like 14 (Drosophila) | 0.0801 |
| 32 | CYAT1 | cyclosporin A transporter 1 | 0.1657 |
| 33 | HLA-DRB1 /// HLA-DRB3 /// HLA-DRB4 /// HLA-DRB5 /// LOC100294036 | major histocompatibility complex, class II, DR beta 1 /// major histocompatibility complex, class II, DR beta 3 /// major histocompatibility complex, class II, DR beta 4 /// major histocompatibility complex, class II, DR beta 5 /// similar to HLA class II histocompatibility antigen, DRB1-7 beta chain | 0.0422 |
| 34 | FLJ10357 | protein SOLO | -0.0966 |

Table S2: *Dengue microarray data: Genes with non-zero weights in the second scPCA loading vector.*

| | Gene Symbol | Gene Name | Weight |
|---|---|---|---|
| 1 | PRSS33 | protease, serine, 33 | 0.1822 |

Table S2: *Dengue microarray data: Genes with non-zero weights in the second scPCA loading vector.*

| | Gene Symbol | Gene Name | Weight |
|---|---|---|---|
| 2 | IFI27 | interferon, alpha-inducible protein 27 | -0.0147 |
| 3 | PI3 | peptidase inhibitor 3, skin-derived | 0.1692 |
| 4 | SLC2A5 | solute carrier family 2 (facilitated glucose/fructose transporter), member 5 | 0.0701 |
| 5 | MYOM2 | myomesin (M-protein) 2, 165kDa | -0.0278 |
| 6 | HLA-DRB4 | major histocompatibility complex, class II, DR beta 4 | -0.0620 |
| 7 | IGH@ /// IGHA1 /// IGHD /// IGHG1 /// IGHG3 /// IGHG4 /// IGHM /// IGHV3-23 /// IGHV4-31 /// LOC100290146 /// LOC100290528 | immunoglobulin heavy locus /// immunoglobulin heavy constant alpha 1 /// immunoglobulin heavy constant delta /// immunoglobulin heavy constant gamma 1 (G1m marker) /// immunoglobulin heavy constant gamma 3 (G3m marker) /// immunoglobulin heavy constant gamma 4 (G4m marker) /// immunoglobulin heavy constant mu /// immunoglobulin heavy variable 3-23 /// immunoglobulin heavy variable 4-31 /// hypothetical protein LOC100290146 /// similar to pre-B lymphocyte gene 2 | 0.4987 |
| 8 | IGKV3-20 | Immunoglobulin kappa variable 3-20 | 0.1623 |
| 9 | RSAD2 | radical S-adenosyl methionine domain containing 2 | -0.2294 |
| 10 | USP18 | ubiquitin specific peptidase 18 | -0.4599 |
| 11 | SIGLEC1 | sialic acid binding Ig-like lectin 1, sialoadhesin | -0.2750 |
| 12 | KCTD14 | potassium channel tetramerisation domain containing 14 | -0.3142 |
| 13 | FAM118A | family with sequence similarity 118, member A | -0.1382 |
| 14 | | | 0.0269 |
| 15 | SLC16A14 | solute carrier family 16, member 14 (monocarboxylic acid transporter 14) | 0.3232 |
| 16 | ANKRD22 | ankyrin repeat domain 22 | -0.2800 |
| 17 | KLC3 | kinesin light chain 3 | 0.1021 |
| 18 | SIGLEC1 | sialic acid binding Ig-like lectin 1, sialoadhesin | -0.0434 |

Table S3: *Dengue microarray data: Gene set enrichment analysis.* The Broad Institute's online gene set enrichment analysis (GSEA) tool was used to identify the ten most significant gene sets based on GO biological processes [13, 9, 10].

| Gene Set Name | Description | Genes in Overlap | *p*-value | FDR *q*-value |
|---|---|---|---|---|
| GO_DEFENSE_RESPONSE | Reactions, triggered in response to the presence of a foreign body or the occurrence of an injury, which result in restriction of damage to the organism attacked or prevention/recovery from the infection caused by the attack. | 13 | 1.12 e-8 | 8.2 e-5 |
| GO_RESPONSE_TO_CYTOKINE | Any process that results in a change in state or activity of a cell or an organism (in terms of movement, secretion, enzyme production, gene expression, etc.) as a result of a cytokine stimulus. | 10 | 3.15 e-7 | 1.16 e-3 |
| GO_MITOTIC_CELL_CYCLE_CHECKPOINT | A cell cycle checkpoint that ensures accurate chromosome replication and segregation by preventing progression through a mitotic cell cycle until conditions are suitable for the cell to proceed to the next stage. | 5 | 8.39 e-7 | 2.06 e-3 |
| GO_CYTOKINE_MEDIATED_SIGNALING_PATHWAY | A series of molecular signals initiated by the binding of a cytokine to a receptor on the surface of a cell, and ending with regulation of a downstream cellular process, e.g. transcription. | 8 | 1.38 e-6 | 2.4 e-3 |
| GO_PROTEIN_LOCALIZATION_TO_CHROMOSOME_ME_CENTROMERIC_REGION | Any process in which a protein is transported to, or maintained at, the centromeric region of a chromosome. | 3 | 1.63 e-6 | 2.4 e-3 |
| GO_CELL_CYCLE_CHECKPOINT | A cell cycle process that controls cell cycle progression by monitoring the integrity of specific cell cycle events. A cell cycle checkpoint begins with detection of deficiencies or defects and ends with signal transduction. | 5 | 3.15 e-6 | 3.86 e-3 |
| GO_POSITIVE_REGULATION_OF_VASOCONSTRICRICTION | Any process that activates or increases the frequency, rate or extent of vasoconstriction. | 3 | 4.74 e-6 | 4.97 e-3 |
| GO_NEGATIVE_REGULATION_OF_METAPHASE_AN_ANAPHASE_TRANSITION_OF_CELL_CYCLE | Any process that stops, prevents or reduces the frequency, rate or extent of metaphase/anaphase transition of cell cycle. | 3 | 8.15 e-6 | 6.58 e-3 |
| GO_MITOTIC_CELL_CYCLE | Progression through the phases of the mitotic cell cycle, the most common eukaryotic cell cycle, which canonically comprises four successive phases called G1, S, G2, and M and includes replication of the genome and the subsequent segregation of chromosomes into daughter cells. In some variant cell cycles nuclear replication or nuclear division may not be followed by cell division, or G1 and G2 phases may be absent. | 8 | 8.6 e-6 | 6.58 e-3 |
| GO_INFLAMMATORY_RESPONSE | The immediate defensive reaction (by vertebrate tissue) to infection or injury caused by chemical or physical agents. The process is characterized by local vasodilation, extravasation of plasma into intercellular spaces and accumulation of white blood cells and macrophages. | 7 | 9.3 e-6 | 6.58 e-3 |

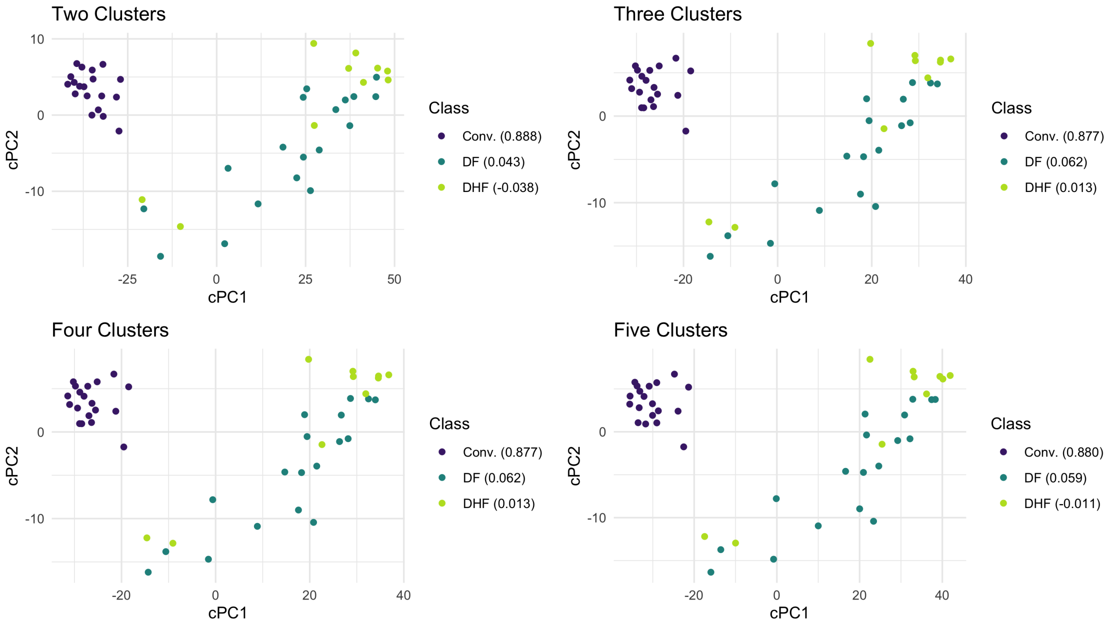


Figure S6: *Dengue microarray data: cPCA.* When varying the *a priori* specified number of clusters for cPCA, all four embeddings are virtually identical, suggesting that cPCA is robust to misspecifications of the number of clusters and that optimal contrastive parameters were selected in each case.

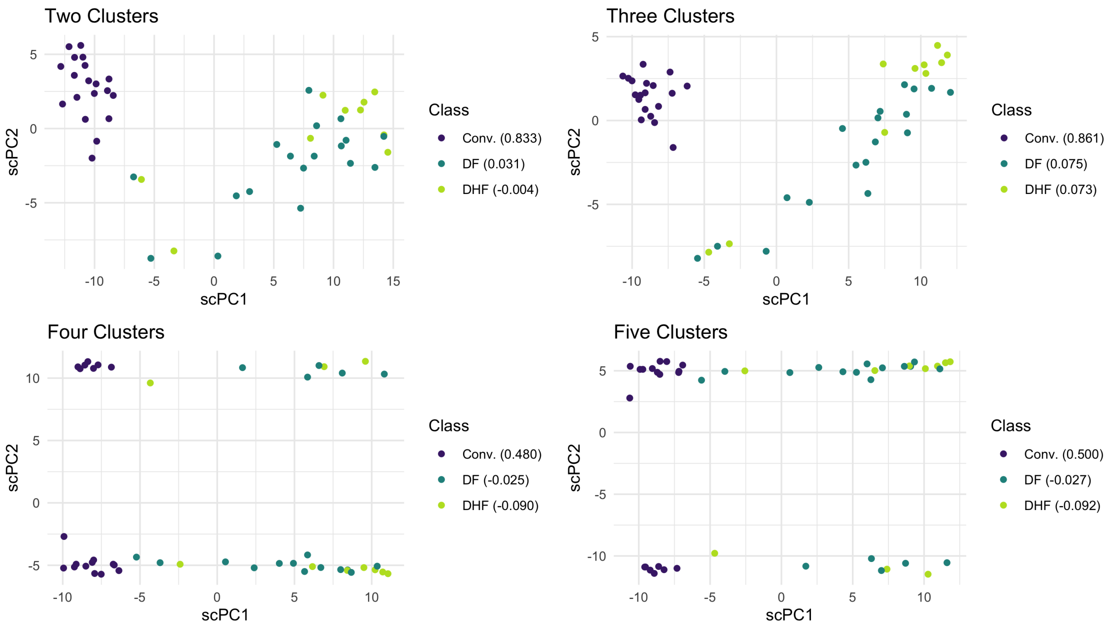


Figure S7: *Dengue microarray data: scPCA.* When varying the *a priori* specified number of clusters for scPCA, we find that the two-dimensional embeddings are sensitive to this choice. When scPCA is performed on this data with four and five clusters, the results resemble those produced by PCA.

#### S6 Leukemia Patient scRNA-seq Data

See Section 3.3 for information on the data. cPCA and scPCA were fit to both patients' data using the non-cross-validated hyperparameter tuning framework. Both methods considered a vector of 40 logarithmically spaced values between 0.1 and 1,000 as potential contrastive parameters. scPCA considered 20 logarithmically spaced values between $1e-9$ and 1 as potential $\ell_1$ penalty parameters. Two t-SNE embeddings were produced per patient: one with an initial dimension reduction step performed with PCA (retaining the 50 leading principal components) and one without. The remaining parameters for each embedding were set to their defaults, except for `theta` which was set to 0. UMAP was performed with its default parameters, except for `n_neighbors` and `min_dist` which were set to 30 and 0.02, respectively. These values match those used by Becht *et al.* [3]. The SIMLR embeddings were produced with `k = 30`, as recommended in Wang *et al.* [15] for datasets of this size. The number of pre-specified clusters was set to 2.

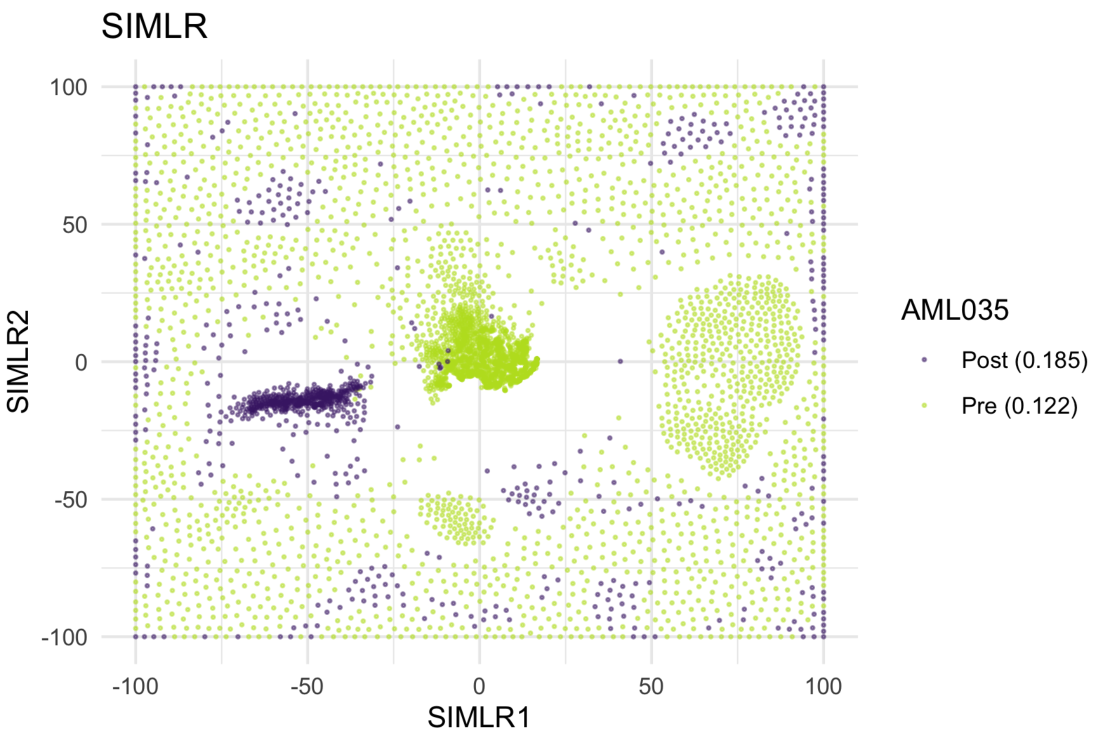


Figure S8: *AML Patient 035 scRNA-seq data: SIMLR.* SIMLR fails to produce an informative two-dimensional embedding of the patient's BMMCs. Although a number of dense clusters are formed in the center of the figure, the spread of observations around these clusters renders the visualization uninterpretable.

Table S4: *AML Patient 035 scRNA-seq data: Gene set enrichment analysis.* The Broad Institute's online gene set enrichment analysis tool was used to identify the ten most significant gene sets based on GO biological processes [13, 9, 10].

| Gene Set Name | Description | Genes in Overlap | $p$-value | FDR $q$-value |
|---|---|---|---|---|
| GO_ESTABLISHMENT_OF_PROTEIN_LOCALIZATIATION_TO_ENDOPLASMIC_RETICULUM | The directed movement of a protein to a specific location in the endoplasmic reticulum. | 33 | 9.82 e-57 | 7.22 e-53 |
| GO_COTRANSLATIONAL_PROTEIN_TARGETING_TG_TO_MEMBRANE | The targeting of proteins to a membrane that occurs during translation. The transport of most secretory proteins, particularly those with more than 100 amino acids, into the endoplasmic reticulum lumen occurs in this manner, as does the import of some proteins into mitochondria. | 32 | 2.09 e-56 | 7.66 e-53 |
| GO_TRANSLATIONAL_INITIATION | The process preceding formation of the peptide bond between the first two amino acids of a protein. This includes the formation of a complex of the ribosome, mRNA or circRNA, and an initiation complex that contains the first aminoacyl-tRNA. | 36 | 1 e-54 | 2.46 e-51 |
| GO_NUCLEAR_TRANSCRIBED_MRNA_CATABOLIC_IC_PROCESS_NONSENSE_MEDIATED_DECAY | The nonsense-mediated decay pathway for nuclear-transcribed mRNAs degrades mRNAs in which an amino-acid codon has changed to a nonsense codon; this prevents the translation of such mRNAs into truncated, and potentially harmful, proteins. | 32 | 4.32 e-54 | 7.94 e-51 |
| GO_PROTEIN_LOCALIZATION_TO_ENDOPLASMICMIC_RETICULUM | A process in which a protein is transported to, or maintained in, a location within the endoplasmic reticulum. | 33 | 1.45 e-53 | 2.13 e-50 |
| GO_VIRAL_GENE_EXPRESSION | A process by which a viral gene is converted into a mature gene product or products (proteins or RNA). This includes viral transcription, processing to produce a mature RNA product, and viral translation. | 34 | 7.2 e-51 | 8.82 e-48 |
| GO_PROTEIN_TARGETING_TO_MEMBRANE | The process of directing proteins towards a membrane, usually using signals contained within the protein. | 34 | 2.71 e-50 | 2.85 e-47 |
| GO_NUCLEAR_TRANSCRIBED_MRNA_CATABOLIC_IC_PROCESS | The chemical reactions and pathways resulting in the breakdown of nuclear-transcribed mRNAs in eukaryotic cells. | 33 | 1.22 e-47 | 1.12 e-44 |
| GO_ESTABLISHMENT_OF_PROTEIN_LOCALIZATIATION_TO_MEMBRANE | The directed movement of a protein to a specific location in a membrane. | 36 | 5.88 e-46 | 4.8 e-43 |
| GO_PROTEIN_TARGETING | The process of targeting specific proteins to particular regions of the cell, typically membrane-bounded subcellular organelles. Usually requires an organelle specific protein sequence motif. | 37 | 2.61 e-43 | 1.92 e-40 |

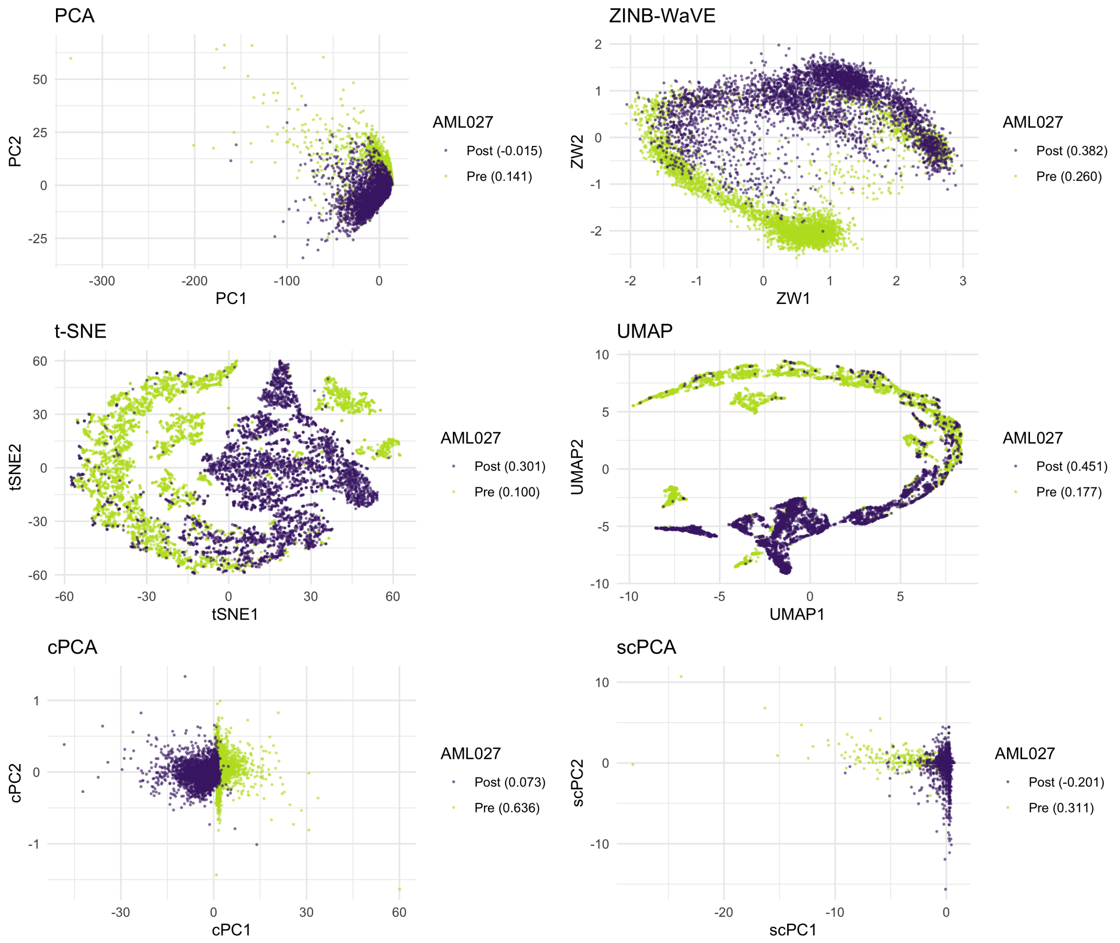


Figure S9: *AML Patient 027 scRNA-seq data.* Two-dimensional embeddings of the patient's BMMCs produced by PCA, ZINB-WaVE, t-SNE, UMAP, cPCA, and scPCA. cPCA and scPCA produce two-dimensional representations that distinguish between the pre- and post-transplant cells of Patient 027. Although cPCA's embedding contains denser clusters, scPCA's clusters are more distinct – though they are oddly shaped. This is the result of sparsity: the scPCA embedding is produced with the count data of only five genes.

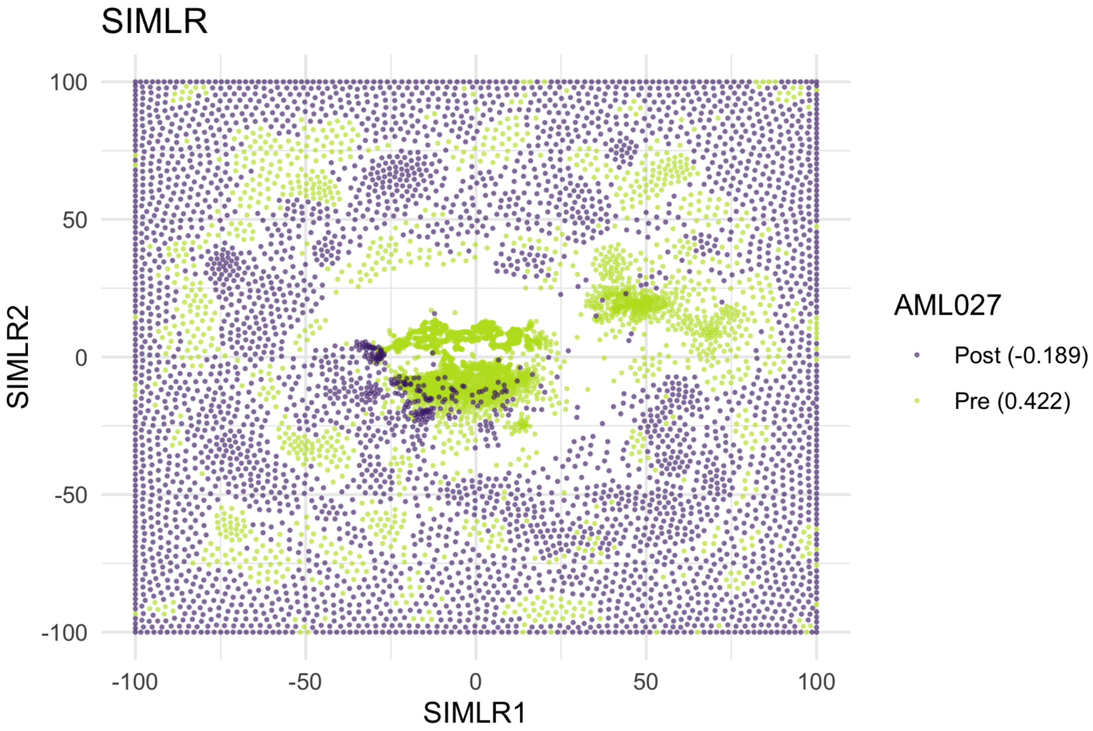


Figure S10: *AML Patient 027 scRNA-seq data: SIMLR.* As with AML Patient 035's data, SIMLR produces an uninterpretable representation of the patient's BMMCs.

#### S7 Mouse Protein Expression Data

Down Syndrome, the leading genetic cause of intellectual disability [7], is the result of trisomy of all or part of the long arm of chromosome 21 [2]. Recently, researchers have begun exploring the use of pharmacotherapies to mitigate these cognitive deficits using the Ts65Dn mouse model [2, 6]. Though not a perfect model for the study of Down Syndrome, the Ts65Dn displays many relevant neurological phenotypic features, such as deficits in learning and memory [12].

Ahmed *et al.* [2] analyzed protein expression in the hippocampus and cortex of Ts65Dn and control mice after exposure to context fear conditioning and Memantine treatment. Memantine, a drug often prescribed to Alzheimer's patients, has been demonstrated to improve performance of the Ts65Dn in tasks that reflect cognitive abilities [2]. The corresponding dataset was made available by Higuera et al. [6]. The data consist of normalized expression measures for 77 proteins from subcellular fractions of the cortex assayed from 38 control and 34 Ts65Dn mice. Each protein expression measurement was repeated 15 times (i.e., 15 technical replicates per mouse for each of the 77 proteins), though a small number of replicates contain missing protein expression measurements due to technical artifacts [6]. More details on the experimental design are provided in Figure S11.

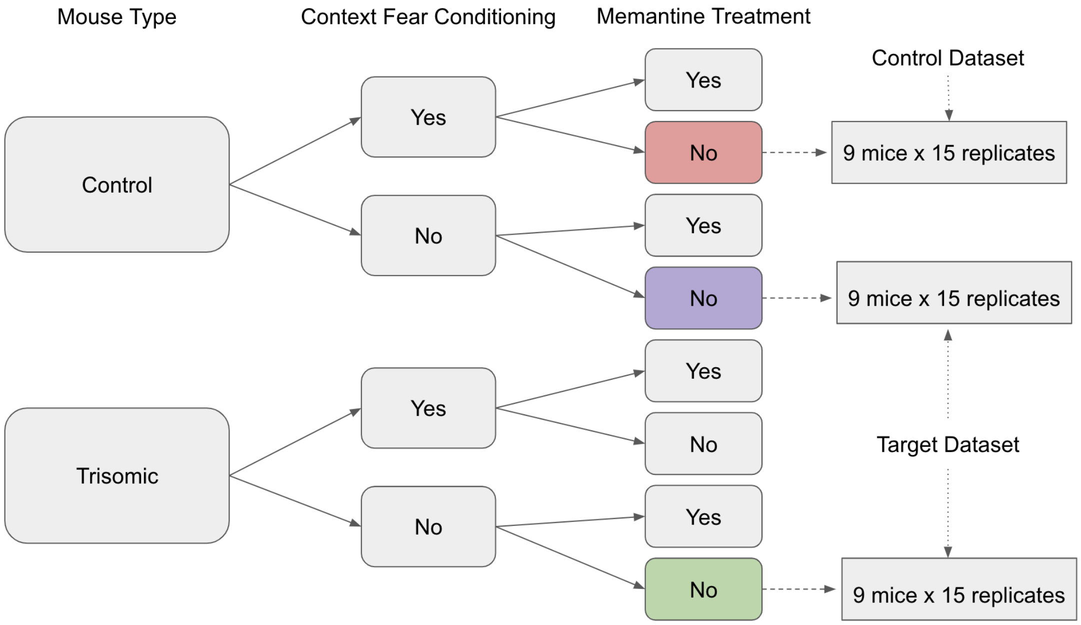


Figure S11: *Mouse protein expression data: Experimental design.* The control dataset is comprised of protein expression measurements for 15 technical replicates from each of 9 control mice subject to context fear conditioning and given a placebo (red leaf). The target dataset consists of protein expression measurements for 15 technical replicates from each of 9 control mice not subject to context fear condition and given a placebo (purple leaf) and 15 technical replicates from each of 9 trisomic mice not subject to context fear condition and given a placebo (green leaf).

To demonstrate scPCA's capacity to capture biologically meaningful and interpretable variation in protein expression data, the technical replicates of the subset comprising 9 control and 9 Ts65Dn mice not subject to context fear conditioning and given a placebo were designated as the target dataset. The technical replicates of the subset of 9 control mice that were subject to context fear conditioning and given a placebo made up the background dataset, as the variation in their protein expression measurements are believed to be similar to that found in the control mice of the target dataset. The data are identical to those used by Abid *et al.* [1] to demonstrate cPCA. PCA, t-SNE, UMAP, cPCA, and scPCA were applied to the target dataset (Figure S12A) to identify differences in protein expression between the control and trisomic mice not exposed to the context fear conditioning experiment. In addition to the target dataset, cPCA and scPCA took as input the column-centered background dataset and specified two clusters *a priori*. cPCA and scPCA were fit using the non-cross-validated hyperparameter tuning framework. Both methods considered a vector of 40 equally spaced values between 0.1 and 1,000 as potential contrastive parameters. scPCA used a vector of 20 equidistant values between 0.05 and 1 as potential $\ell_1$ penalty parameters. t-SNE embeddings were produced with and without an initial dimensionality reduction step step. The remaining variables were left at their defaults, with the exception of `theta` which was set to 0. The embedding produced without an initial application of PCA produced the best embedding, and so it is presented in the manuscript and supplement. The UMAP embedding was generated with `n_neighbors` set to 30 and with `min_dist` set to 0.02. The remaining parameters were set to their defaults.

PCA proved incapable of distinguishing between the biological groups of interest. UMAP, cPCA, and scPCA successfully split the control and trisomic mice into virtually distinct clusters, though the number of clusters found by UMAP and cPCA in two dimensions did not match, even when varying the *a priori* specified number of clusters in cPCA (Figure S14). Comparing the results of UMAP and scPCA, we find that they produce the same number of clusters, but their representations of the global structure are markedly different, even when varying the number of clusters specified *a priori* in scPCA (Figure S15). The presence of distinct Ts65Dn clusters in UMAP's representation may correspond to technical noise that is diminished in cPCA's and scPCA's embeddings, or may arise from UMAP's inability to dependably capture global structure. We also remark that cPCA and scPCA produce very similar embeddings, up to a rotation; however, the first and second columns of scPCA's loading matrix contain merely 12 and 16 non-zero entries, respectively (Figure S12B). Also note that the the separation of control and trisomic mice by scPCA only occurs in scPC2: the proteins with non-zero weights in the corresponding loading vector include AKT, APP, SOD1, and GSK3, each of which has been associated with Down Syndrome in human or mouse models [14, 11, 5, 8]. The full list of proteins with non-zero weights in the first two loading vectors of scPCA are provided in Table S5 and Table S6.

Table S5: *Mouse protein expression data: Proteins with non-zero weights in the first scPCA loading vector.*

| | Protein Symbol | Weight |
|---|---|---|
| 1 | ELK | 0.0618 |
| 2 | BRAF | -0.1001 |
| 3 | RSK | -0.0927 |
| 4 | SOD1 | 0.1800 |
| 5 | S6 | 0.1281 |
| 6 | AcetylH3K9 | 0.3992 |
| 7 | RRP1 | 0.0606 |
| 8 | Tau | 0.7320 |
| 9 | CASP9 | -0.0795 |
| 10 | PSD95 | -0.0329 |
| 11 | Ubiquitin | -0.3674 |
| 12 | H3AcK18 | 0.2958 |

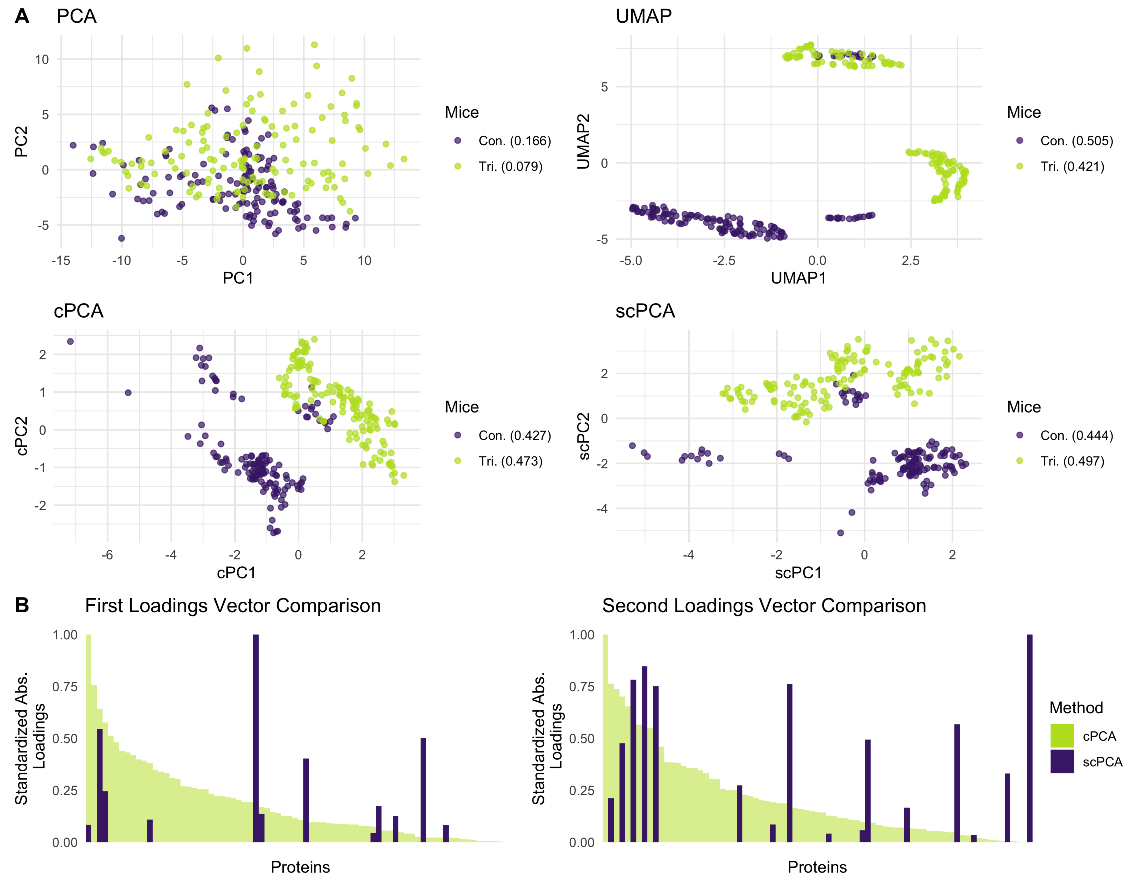


Figure S12: *Mouse protein expression data.* **A** All methods but PCA were capable of separating the control from the trisomic mice, though it is unclear why UMAP splits the Ts65Dn mice into two distinct groups. scPCA's low-dimensional representation of the protein expression data is markedly similar to that of cPCA, up to a rotation, despite relying on only a fraction of non-zero values in the loading matrix. On average, scPCA also produces the tightest clusters. Note: a small group of control mice were clustered with the trisomic mice in the UMAP, cPCA, and scPCA representation, potentially comprising a group of mislabeled mice. **B** scPCA's leading vectors of loadings are much sparser than those of cPCA, increasing the interpretability of findings and clarity of the visualization. The differing rotations of cPCA and scPCA, in addition to the drastically different weighting scheme of the proteins in their respective loading matrices, may indicate that the contrastive step performed by cPCA does not sufficiently dampen spurious sources of variation in the data.

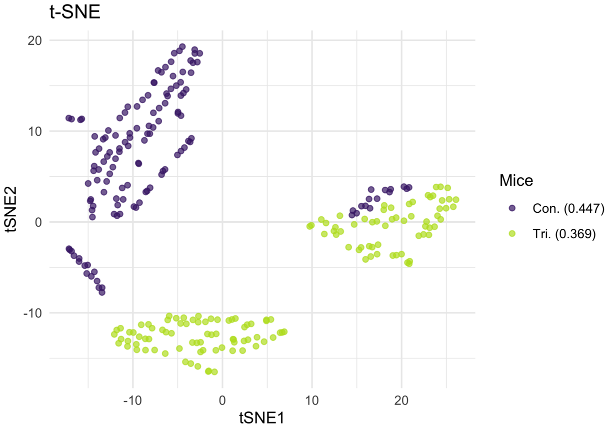


Figure S13: *Mouse protein expression data: t-SNE.* t-SNE produces almost linearly-separable clusters, though these clusters contain many fractured, spurious sub-clusters that do not relate to biological signal.

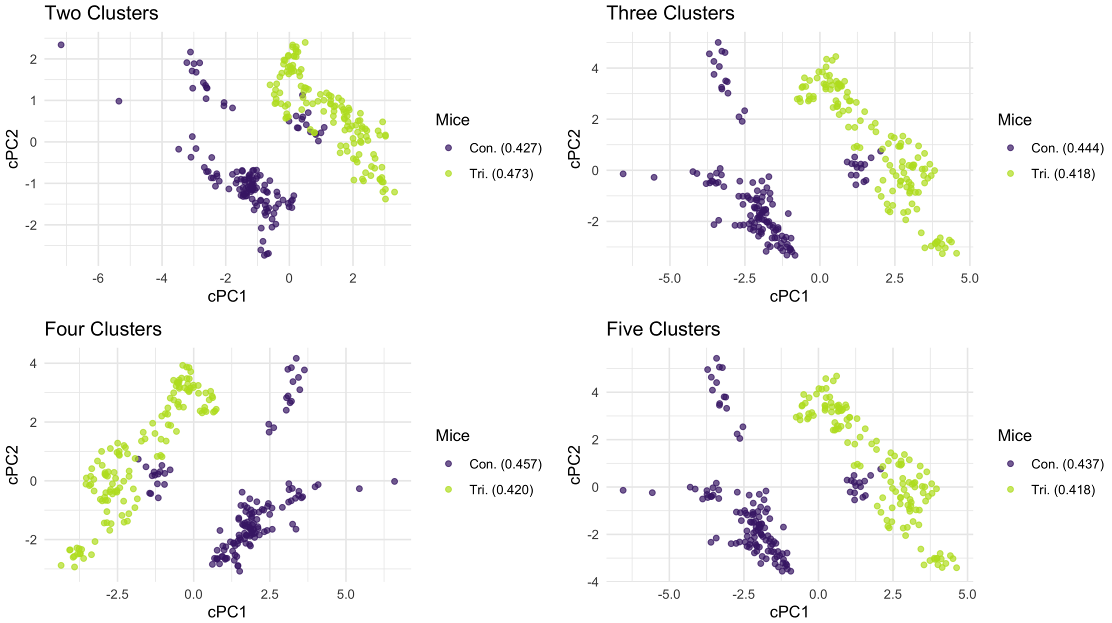


Figure S14: *Mouse protein expression data: cPCA.* When varying the *a priori* specified number of clusters for cPCA, we find that the two-dimensional embedding is once again robust to misspecifications.

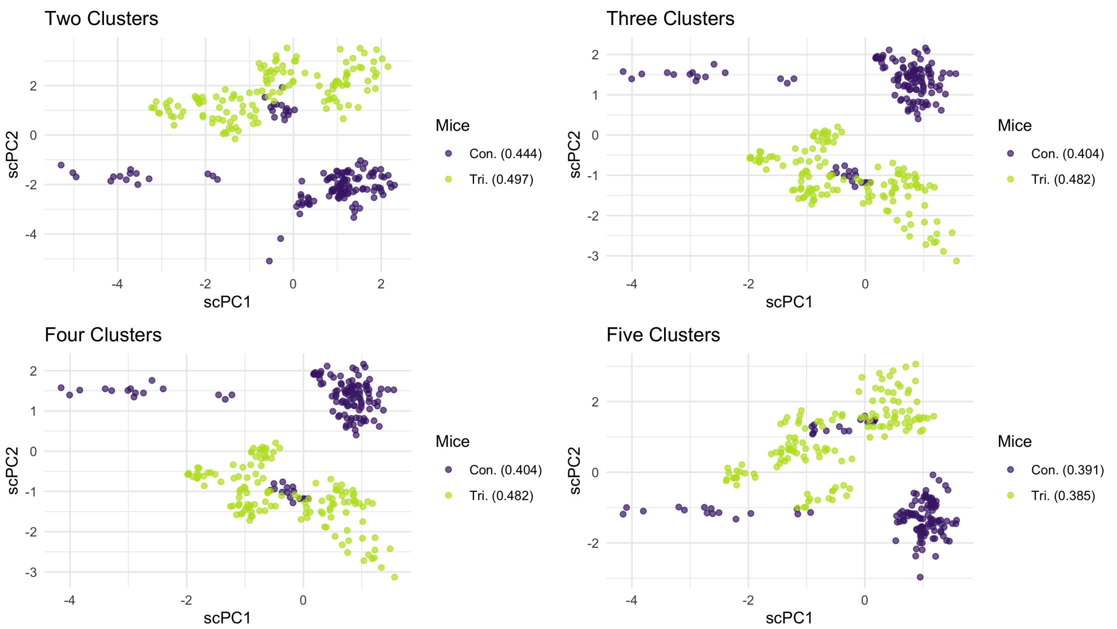


Figure S15: *Mouse protein expression data: scPCA.* Unlike with the dengue microarray data, when varying the *a priori* specified number of clusters for scPCA, we find that the two-dimensional embedding is robust to misspecifications. This may indicate that the sensitivity of the method to this tuning parameter is data-dependent.

Table S6: *Mouse protien expression data: Proteins with non-zero weights in the second scPCA loading vector.*

| | Protein Symbol | Weight |
|---|---|---|
| 1 | ELK | -0.2236 |
| 2 | AKT | -0.0999 |
| 3 | APP | 0.3525 |
| 4 | SOD1 | -0.2323 |
| 5 | NUMB | 0.4690 |
| 6 | P70S6 | 0.1554 |
| 7 | GSK3B | 0.3574 |
| 8 | PKCG | 0.3978 |
| 9 | S6 | 0.3674 |
| 10 | RRP1 | 0.0272 |
| 11 | GluR4 | 0.0404 |
| 12 | IL1B | -0.2664 |
| 13 | P3525 | 0.0196 |
| 14 | PSD95 | -0.1289 |
| 15 | SNCA | -0.0783 |
| 16 | H3AcK18 | 0.0169 |

#### S8 Cross-Validated cPCA and scPCA

The cross-validated (CV) versions of cPCA and scPCA (as implemented in Algorithm 2) were applied to the datasets presented in the main paper and in the supplement. The hyperparameter grids used by each method are the same as those employed for their non-cross-validated counterparts. See Sections S4, S5, S6, and S7 for details. Five-fold CV was applied to the simulated scRNA-seq data, mouse protein expression data, and AML patient scRNA-seq data (Figures S16, S17, S19, and S20), and 3-fold CV was applied to the dengue microarray gene expression data (Figure S18). A reduced number of folds was used on the latter dataset since it possessed much fewer observations than the others. For each dataset, the CV-cPCA and CV-scPCA embeddings closely resemble – and are in some cases identical to – their non-cross-validated counterparts.

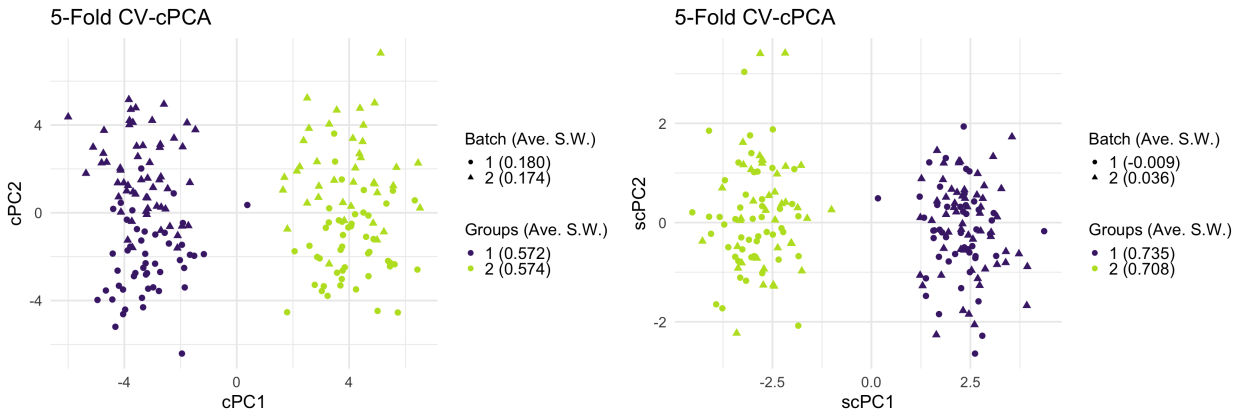


Figure S16: *CV-cPCA and CV-scPCA on simulated scRNA-seq data.*

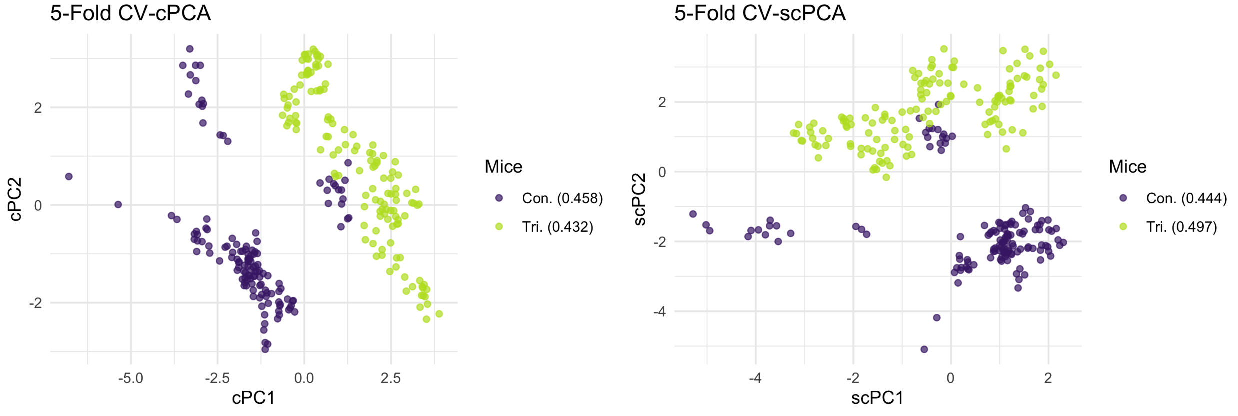


Figure S17: *CV-cPCA and CV-scPCA on mouse protein expression data.*

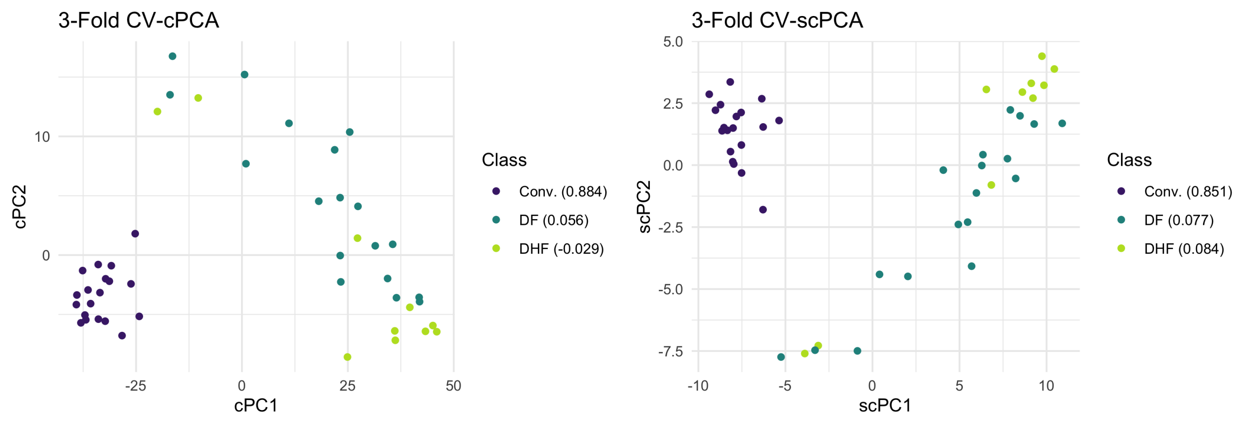


Figure S18: *CV-cPCA and CV-scPCA on dengue microarray data.*

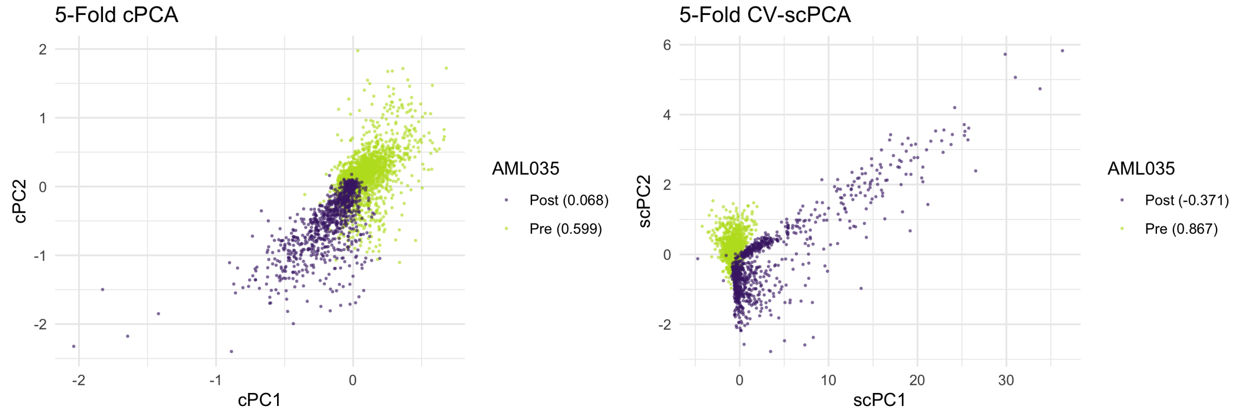


Figure S19: *CV-cPCA and CV-scPCA on AML Patient 035 scRNA-seq data.*

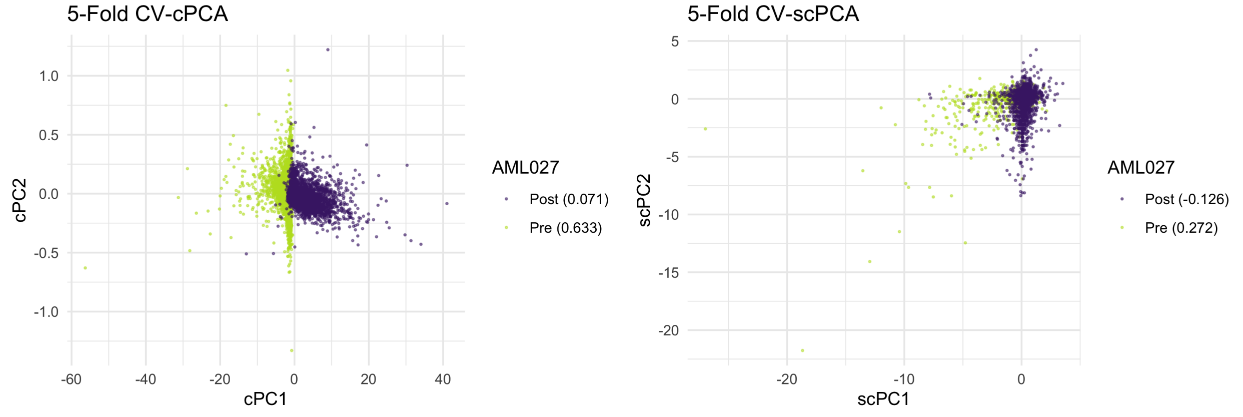


Figure S20: *CV-cPCA and CV-scPCA on AML Patient 027 scRNA-seq data*

#### S9 Running Time Comparison

The running times of PCA, t-SNE, UMAP, cPCA, scPCA, ZINB-WaVE, and SIMLR were recorded on the simulated scRNA-seq data from Section 3.1 and on the scRNA-seq data of AML Patient 035 from Section

3.3. If a method's software implementation offered the option to parallelize, four cores were used. The hyperparameters used by the methods on each dataset are identical to those described in their respective sections of the supplement, that is Sections S4 and S6. The `microbenchmark R` package was used to track the methods' running times. The comparison is presented in Figures S21 and S22.

We note that the iterative algorithm presented in Section 2 is computationally inefficient. However, the scPCA framework is not dependent on Zou *et al.* [16]'s optimization procedure; other solutions to the SPCA criterion (Equation (3)) can be employed to sparsify the loadings of contrastive covariance matrices. Indeed, more efficient algorithms exist.

In particular, recent work by Erichson *et al.* [4] provides a scalable algorithm by reformulating the SPCA criterion (Equation (3)) as a value function optimization problem. Instead of requiring an iterative routine to update $\mathbf{B}$ of the alternating algorithm presented in Section 2, a single operator is used. This procedure can be sped up further through the use of randomized linear algebra methods to compute $\mathbf{A}$.

These sparsification methods were recently included in the `scPCA R` package, and the distribution of their running times are included in Figures S21 and S22. Using these recently developed methods to solve the SPCA step, the scPCA algorithm's running time is decreased by over an order of magnitude on both datasets. Its running time is similar to that of competing methods when using four cores. Given that the hyperparameter tuning framework is embarrassingly parallel, one can expect even faster computation times when more cores are employed.

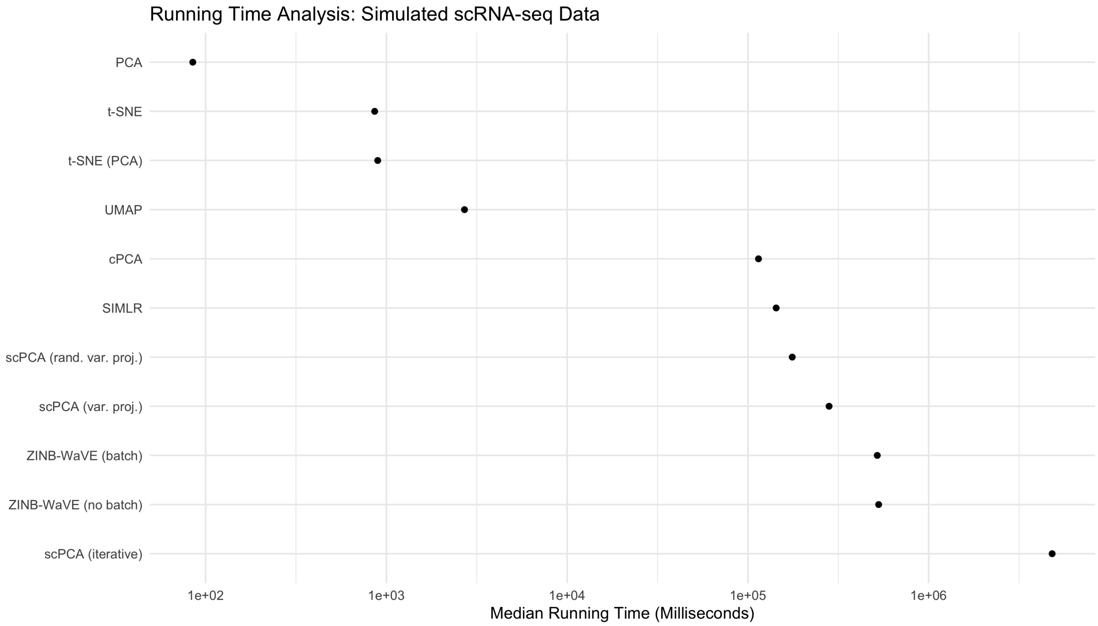


Figure S21: *Running time comparison: Simulated scRNA-seq data.* Each method was applied to the data five times. Note that *scPCA (var. proj.)* and *scPCA (rand. var. proj.)* correspond to the scPCA algorithms relying on the SPCA procedures detailed in Erichson *et al.* [4]'s recent work, the latter being being the procedure which relies on randomized techniques. *scPCA (iterative)* pertains to the scPCA method that uses the SPCA algorithm detailed in Zou *et al.* [16]. The median running time of each method is reported. The most general of the dimensionality reduction methods, PCA, t-SNE, and UMAP, were at least an order of magnitude faster than all other methods. The remaining method's running times were similar, with the exception of the much slower *scPCA (iterative)*.

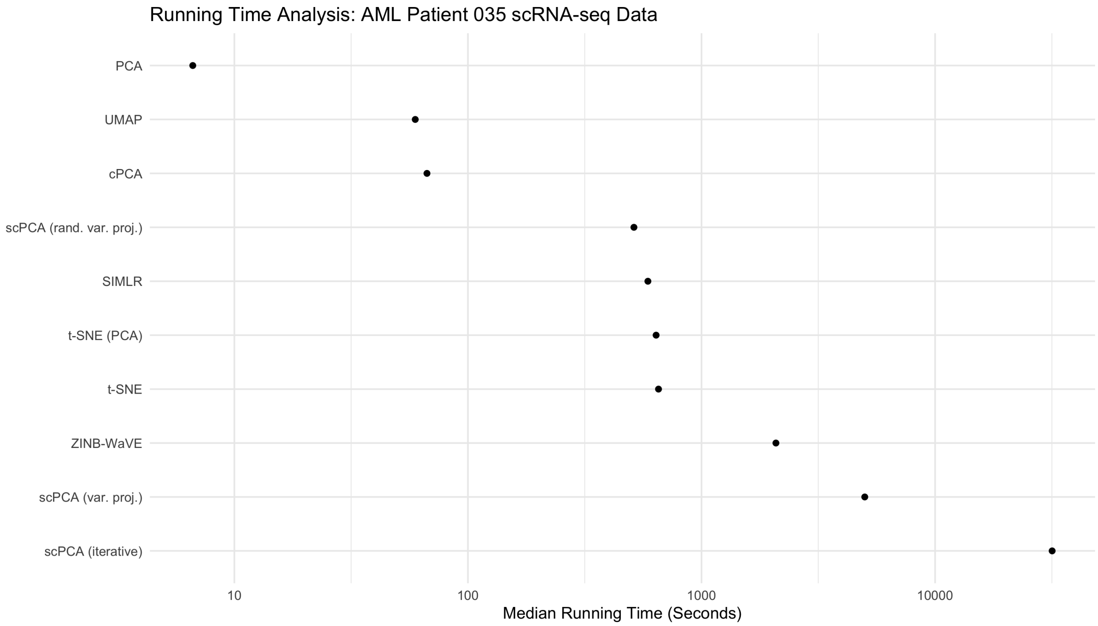


Figure S22: *Running time comparison: AML Patient 035 scRNA-seq data.* Each method was applied to the data three times. The median running time of each method is reported. Compared to the smaller dataset, the contrastive methods presented in the manuscript are competitive with the more general dimensionality reduction methods. Indeed, cPCA's running time is similar to that of UMAP, and the scPCA algorithm relying on random numerical methods for sparsification is faster than t-SNE.
